# Paradigm shift in eukaryotic biocrystallization

**DOI:** 10.1101/2022.01.11.475817

**Authors:** Jana Pilátová, Tomáš Pánek, Miroslav Oborník, Ivan Čepička, Peter Mojzeš

## Abstract

Despite the widespread occurrence of crystalline inclusions in unicellular eukaryotes, scant attention has been paid to their composition, functions, and evolutionary origins, assuming just their inorganic contents. The advent of Raman microscopy, still scarcely used for biological samples, allowed chemical characterization of cellular inclusions *in vivo.* Using this method, herein we provide a substantial revision of the cellular crystalline inclusions across the broad diversity of eukaryotes examining all major supergroups. Surprisingly, here we show that 80 % of these crystalline inclusions contain purines, mostly anhydrous guanine (62 %), guanine monohydrate (2 %), uric acid (12 %) and xanthine (4 %). Hence, our findings indicate that purine biocrystallization is a very general and an ancestral eukaryotic process operating by an as-yet-unknown mechanism. Purine crystalline inclusions are high-capacity and rapid-turnover reserves of nitrogen of a great metabolic importance, as well as optically active elements, *e.g.*, present in the light sensing eyespots of flagellates, possessing even more hypothetical functions. Thus, we anticipate our work to be a starting point for more in-depth studies of this phenomenon on the detailed level spanning from cell biology to global ecology, with further potential applications in biotechnologies, bio-optics or in human medicine.

---

Crystalline inclusions, conspicuous in many single-celled eukaryotes, have attracted the attention of scientists since the emergence of microscopy. After Charles Darwin documented the crystal-like particles in microscopic flagellates^1^, Ernst Haeckel coined the term “biocrystal” for calcite (CaCO_3_), celestite (SrSO_4_), silica (SiO_2_), and oxalate in protists^2^. Biocrystals or biogenically originated crystalline inclusions of either organic or inorganic chemical nature, are typically formed inside vacuoles and grow into shells and scales on cell surfaces, *e.g.*, calcite scales of coccolithophores that take part in the global carbon cycle^3^. On the other hand, rarely-reported purine inclusions of guanine, xanthine, hypoxanthine, or uric acid are often overlooked^4–8^. The recent resurgence of research on purine inclusions has shown rapid uptake kinetics of different nitrogen compounds that are converted into massive guanine nitrogen stores with sufficient capacity, in some cases, to support three consecutive cell cycles^8^. As a fundamental biogenic element, nitrogen represents a great share of biotic elemental stoichiometry, from cellular to global scales, impacting Earth’s climate^9^. Biocrystalized guanine polarizes and reflects light in many animals, *e.g.*, fish with the opalescent guanine crystals in the iridocytes of their scales resembling glittery camouflage, or as an adaptation for vision in scallops, deep-sea fishes and arthropods that exhibit guanine-reflective retinal tapeta in their eyes^10, 11^. Analogous functions of guanine crystals in alveolate protists have also been reported^5, 12, 13^.

To date, the investigation of crystalline inclusions and their biological importance in unicellular eukaryotes has been impeded by a lack of methods for their non-destructive *in situ* and *in vivo* characterization. Although Raman microscopy^8^ provides just such capabilities (as detailed in Supplementary Materials and Methods), this methodology has not yet been fully exploited in the study of intracellular crystalline inclusions. Compared to other tools for microscopic elemental analysis (nanoSIMS, EELS, EDX TEM), or chemical analytical approaches (GC and LC MS, NMR, X-ray crystallography), Raman microscopy offers direct visualization of molecular profiles for various cellular compartments in single cells with the resolution of confocal microscopy and without the need for fixation, time-consuming and laborious sample preparation, or the need for large amounts of biological material necessary for chemical extractions. Thus, it is also suitable for environmental samples. Herein we take a major step forward in the research of crystalline cell inclusions by revisiting their chemical nature across the eukaryotic tree of life using Raman microscopy.

## Results

We have screened all major eukaryotic groups, using >200 species, most of them for the first time, searching for birefringent (light-polarizing) crystalline inclusions. They are commonly located inside vacuoles, wobbling by Brownian motion (movies S1 to S8). Raman spectra of the biocrystals found are displayed in Figs. S1 to S5. We registered common biocrystals in accordance with previously-described crystalline inclusions, such as calcite, oxalate, celestite, and baryte^3^, together with other birefringent structures (*i.e.*, starch, chrysolaminarin, strontianite, and newly observed crystals of sterols and carotenoids). However, apart from these, we found a surprisingly broad occurrence of purines (>80 % of examined species containing crystals), particularly crystalline anhydrous guanine (62 %), uric acid (12 %), xanthine (4 %) and guanine monohydrate (2 %), (Fig. 1; Table S1; Supplementary Text). For the first time, we found anhydrous guanine purine crystal inclusions predominated in model and biotechnologically important species, and also in environmentally important strains. It commonly occurs in cosmopolitan marine and freshwater algae – including bloom-causing dinoflagellates, in the endosymbionts of corals – crucial for the maintenance of entire coral reef ecosystems, in unicellular parasites of warm-blooded animals and in cellulose-digesting anaerobic symbionts of termites, in slime molds etc. The lack of purine inclusions in some strains in our dataset (Table S1) does not exclude the existence of the necessary synthetic pathways since the induction of crystal formation may occur only under specific conditions.

**Fig. 1:**
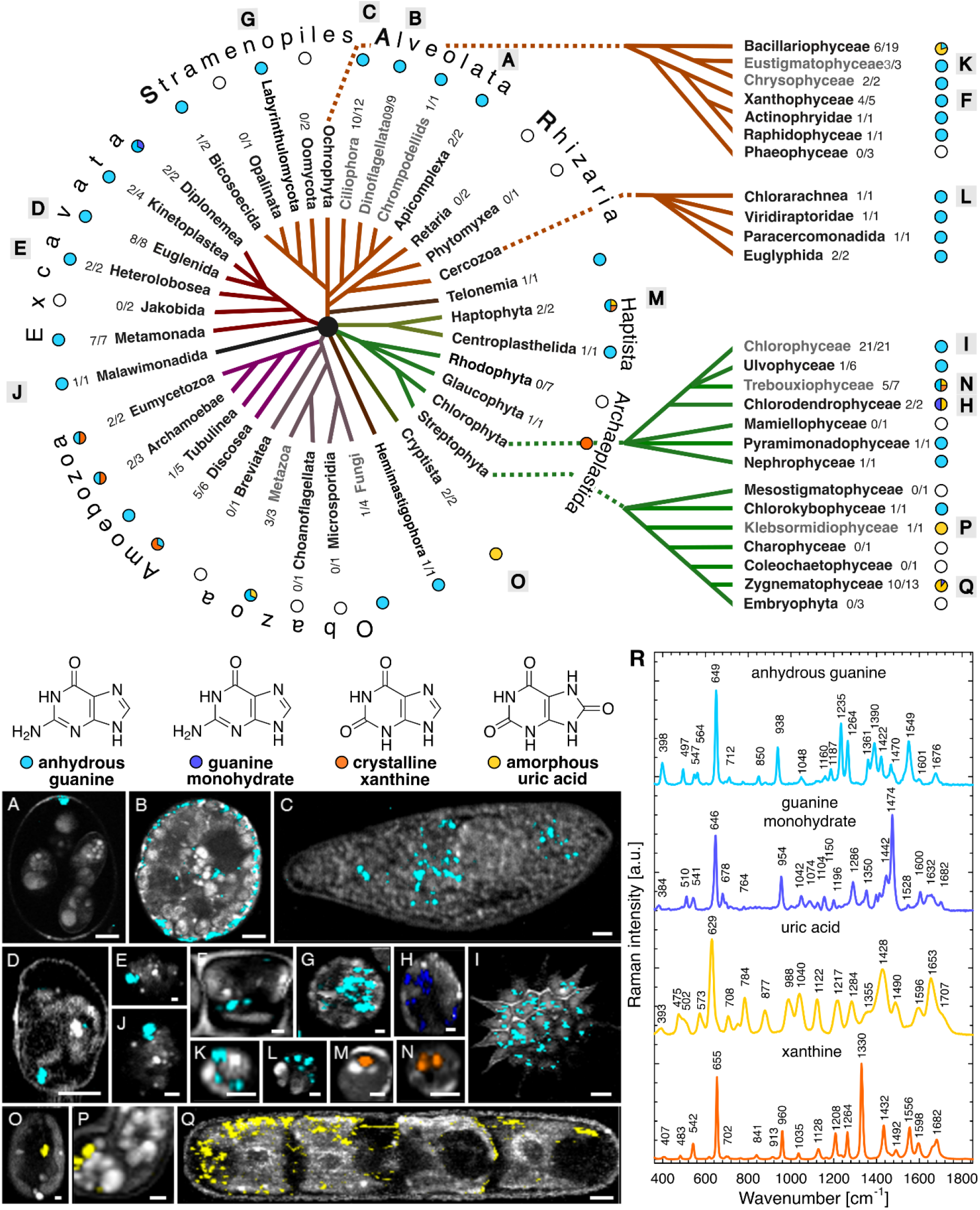
Distribution of purine inclusions identified by Raman microscopy in the eukaryotic tree of life. The occurrence of anhydrous guanine (cyan), guanine monohydrate (violet), uric acid (yellow) and xanthine (orange) are illustrated in the evolutionary scheme as pie charts and in Raman maps (A–Q) together with the Raman spectra (R). Ratio of species positively tested for purine inclusions out of the total number of screened samples are expressed for each taxonomic category. Lineages highlighted in grey possess purine inclusions already reported elsewhere. A – *Eimeria maxima*, B – *Glenodinium foliaceum*, C – *Paramecium* sp., D – *Eutreptiella gymnastica*, E – *Naegleria gruberi*, F – *Tribonema aequale*, G – *Schizochytrium* sp., H – *Tetraselmis subcordiformis*, I – *Pediastrum duplex*, J – *Gefionella okellyi*, K – *Nannochloropsis oculata*, L – *Bigelowiella natans*, M – *Isochrysis* sp., N – *Chlorella vulgaris*, O – *Cryptomonas* sp., P – *Klebsormidium flaccidum*, Q – *Penium margaritaceum* Scale bars: 5µm (A–D, I, Q), 1 µm (E–H).

We detected anhydrous guanine as the constituent of most purine crystals in all of the eukaryotic major groups (Fig. 1). Additionally, we found uric acid in cryptophytes, some diatoms, zygnematophytes, and klebsormidiophytes. We identified xanthine crystals in Amoebozoa, as well as in biotechnologically important microalgae (*e.g.*, *Chlorella* and *Isochrysis*). Interestingly, we discovered the first occurrence of pure guanine monohydrate in marine diplonemids and as an admixture to uric acid in two green algae. Particular triggers for purine inclusion formation are unknown, but they are often produced after transfer to fresh growth media, media containing surplus sources of nitrogen, and under stress conditions^4, 6^. As previously shown, purine crystals may act as nitrogen storage for microalgae in which they are formed in a type of luxury uptake resulting in net removal of nitrogen from the medium^8^.

Intracellular biocrystals, typically occurring inside membrane-bound compartments, are eukaryote-specific with the exception of bacterial magnetosomes^3^. The widespread distribution of purine crystals may be contingent on the emergence of cell compartmentalization in early eukaryotes. Furthermore, our results suggest that purine crystals might have been present in the last eukaryotic common ancestor (LECA), becoming one of the very first types of biocrystals in eukaryotes. Therefore, we employed comparative transcriptomics and genomics to identify candidate proteins that could be involved in the probably ancient pathways responsible for purine crystal formation. The components of the salvage pathway including phosphoribosyltransferases are such plausible candidate proteins. Consistent with this idea, we proved hypoxanthine-guanine phosphoribosyltransferase (HGPT) to be omnipresent among eukaryotes (Fig. S8 and S12) and we hypothesize that it might be responsible for purine reusage after crystal degradation and/or hydrolysis of nucleotides, releasing simple purines in order to form purine crystals. Furthermore, nucleobases (*i.e.,* purines), nucleosides and/or nucleotides must be transported from cytoplasm to the vesicles where crystals are formed. Firstly, we focused on the most straightforward group of purine transporters that are considered to be commonly present among eukaryotes^19^. Our exhaustive homolog search and subsequent phylogenetic analyses surprisingly challenged the ubiquity of the three of them: nucleobase-cation symporter 1 – NCS1, nucleobase-ascorbate transporter – NAT, and AzgA (see Supplementary Text for details). Of these, only the AzgA transporter was probably present in LECA (see Fig. S8 and S9), although distribution of both its eukaryotic paralogs is rather limited. The evolutionary history of the other two known purine transporters in eukaryotes is much more complex and we do not have any strong evidence for their presence in LECA (Fig. S8, S10, and S11). NCS1 exists in eukaryotes in several paralogs, at least three of them emerged by relatively recent horizontal gene transfer from eubacteria (Fig. S10). NAT (= NCS2) emerged independently four times in eukaryotes (Fig. S11). Our final argument against involvement of solo-purine transporters in crystal formation is their absence in some of the purine crystal-forming groups (Heterolobosea, Ciliophora, and Apicomplexa). Next, we analyzed the distribution of nucleoside transporters, *e.g.,* the concentrative (CNT) and equilibrative (ENT) nucleoside transporters, showing that CNT has an infrequent occurrence in eukaryotes (Table S2). The ENT family appears to be the most promising, as it is the only one among all nucleotide/nucleoside/nucleobase transporters we tested that has a clear pan-eukaryotic distribution (Table S3). Members of the ENT family are specific for nucleosides and nucleobases, and are part of the major facilitator superfamily (MFS). In general, ENTs can operate in a bidirectional mode, in some cases with cation symport and with different localization in the plasma membrane or in intracellular vesicles^20^. There are also other candidate carriers (*e.g.*, VNUT that is known from metazoans), but the exact distribution, function, and localization of such proteins cannot be reliably predicted *in silico* without further biochemical studies in other eukaryotes^21^. The metabolism of nucleobases, nucleosides and nucleotides together with their transport is essential for all organisms and hence there may not be any purine transporters solely involved in crystal formation, as they likely play additional essential roles in the cell. Thus, extensive biochemical and proteomic studies have to be employed to answer this question in future.

## Discussion

Intriguingly, the transition of crystal composition from purines in green algae to calcium oxalate in land plants may be metabolically bridged through purine degradation^14, 15^. We also see a similar trend in Fungi, with yeasts possessing purine crystals, filamentous fungi producing calcium oxalate^7, 16^ and some marine seaweeds (Phaeophyceae) lacking crystals altogether. Hence, loss of the capacity to form purine crystals may correlate with the development of multicellularity that the necessity to store nitrogen is replaced by transfer of soluble metabolites through the multicellular body^17^. In some animals purine crystals, rather than serving as metabolic depots, instead function as photonic mirrors^10, 11^. Others produce purines as a waste product mitigating the toxicity of excessive nitrogen uptake when excretion is not possible or is outstripped^18^. However, nitrogen is a growth-limiting factor for free-living microscopic eukaryotes^9^, and thus, a mechanism to store this essential element instead of wasting it is a significant advantage. Due to low-solubility and high-capacity, nitrogen-rich purine inclusions might have emerged as a preadaptation to nitrogen detoxification, protecting against exposure to high levels of ammonia or nitrates, utilizing vacuoles as a versatile sequestration space. On the other hand, in nitrogen depleted environments, nitrogen storage may mitigate stress-induced oxidative by allowing prompt production of glutathione, heat-shock proteins, chaperones, peroxidases, etc.

Raising awareness of the existence of purine inclusions in diverse unicellular eukaryotes brings implications for other fields as well. The nitrogen-rich microbes might be of use in biofertilizers. The exceptional optical activity of purine crystals can be exploited in cosmetics for pearly iridescent effects or in optics for scalable surfaces with magnetically alterable reflectivity^10^. The value of algae-based food supplements may be limited by the medical issues associated with regular intake, of purines, *e.g.*, hyperuricemia manifesting as gouty arthritis and increased cardiovascular risk, related to accelerated atherosclerosis^22^. Additionally, producers of algae-based biofuels and nutraceuticals commonly use nitrogen starvation to stimulate lipid production^23^. The implications of purine inclusions associated with this process have not yet been taken into account.

To understand the process of purine crystals formation and degradation may be crucial for treatments of hyperuricemia-caused urolithiasis or gout in human medicine. According to epidemiological evidence, the increasing prevalence of hyperuricemia (∼21 %) as one of the modern civilization diseases represents the main cause of gout^22^. Gout and uric acid nephropathy are caused by the purine crystals formation in joints and kidneys, respectively, and they are still of unknown mechanism. The current treatment is limited to xanthine oxidase inhibitors (*e.g.*, allopurinol), as in most of the cases, there is no way to dissolve or remove the crystals already formed^22^. In the course of evolution, humans might have lost the ability for purine crystals degradation. Thus, the understanding of purine biocrystallization in unicellular models can help to understand human pathophysiology and the molecular traits of both: purine crystal formation in order to design specific inhibitors, as well as the mechanism of degradation in unicellular organism in order to introduce this system to the multicellular bodies for establishing the curative treatments of hyperuricemia-related diseases.

Purine inclusions, often overlooked but widely distributed in unicellular eukaryotes, serve as a high-capacity nitrogen storage mechanism and comprise a previously missing piece of the unsolved puzzle of the global nitrogen cycle. Indeed, most of the light-polarizing crystals present in various species of eukaryotes appear to contain purines, including guanine, uric acid and xanthine. This is in stark contrast to the traditional assumption that biocrystals are typically formed by the inorganic crystals, calcite and calcium oxalate^2^. The under-appreciated but widespread occurrence of purine inclusions, especially in unicellular eukaryotes, offers a rich source of material for future study in fields from cell biology to global ecology.

## Methods

### Biological material – cell cultures and environmental samples

To assess the composition of various birefringent cell inclusions among microscopic eukaryotes, we screened species from environmental samples, cell cultures obtained from various culture collections, and strains from private collections that were kindly donated by our collaborators (listed in Table S1 and acknowledgements). The cultivation conditions for each of the tested species are shown in Table S1. Environmental samples were assessed within one week after collection. Cell cultures were observed after transfer to fresh media on the same day and/or during five consecutive days until the purine inclusions were observed. In some cases (marked with “*” in Table S1), we transferred the cells to media containing dissolved guanine (approximately 30 µM final concentration), in order to facilitate the formation of crystalline inclusions.

### Polarization microscopy

We used polarization microscopy to screen organisms for intracellular crystalline inclusions. Initially, polarization microscopy was performed separately, using an Olympus AX 70 Provis microscope (Olympus, Japan) or confocal microscope (Leica TCS SP8; Leica, Germany) equipped with a digital camera (Leica MC170 HD; Leica, Germany). After installing the polarization filters directly on the Raman microscope (WITec alpha300 RSA, WITec, Germany), photomicrographs and videos were taken immediately before Raman measurements. Short videos were taken using the default settings (25 fps using the Leica TCS SP8 or 20 fps using the WITec alpha300 RSA). Videos were processed using Vegas Pro 14.0 software (MAGIX, Germany).

### Raman microscopy

Measurements and data-processing were performed as described elsewhere^6, 8, 24^. The advantages and possible drawbacks of this method have recently been discussed^25^. In brief, Raman map scanning of 203 species comprising more than 3 000 measurements of whole cells, and/or single spectra of crystalline inclusions of mobile cells, or cells with fast-moving cytoplasm, was done using a confocal Raman microscope (WITec alpha300 RSA) equipped with the following objectives: 20× EC Epiplan, NA = 0.4 (Zeiss, Germany), 50× EC Epiplan-Neofluar, NA = 0.55 (Zeiss, Germany), 60× water-immersion UPlanSApo, NA 1.2 (Olympus, Japan), 100× oil-immersion UPlanFLN, NA 1.3 (Olympus, Japan). A 532 nm laser with a power of approximately 20 mW at the focal plane was used.

For Raman map scanning, cell cultures were used as follows: 1 ml of culture was centrifuged at 2000 g for 1 min, when necessary for fast-moving flagellates, immobilization was accomplished by mixing 5 μl of the cell pellet with 5 μl of 1% low-temperature-melting agarose spread under a 20 mm diameter, 0.18mm-thick quartz coverslip sealed with CoverGrip (Biotium, USA). In cases of environmental samples, where assessment of cell movement is crucial for reliable identification of species (mostly Amoebozoa and Excavata), immobilization was not used, and data acquisition was performed *via* Raman single-spectrum mode with an integration time of 0.5 s and 20 accumulations using one of the following objectives: 50× EC Epiplan-Neofluar, 60× UPlanSApo, and 100× UPlanFLN. Raman map measurements were performed with a scanning step of 200 nm in the both directions, voxel size 1 μm^3^ and an integration time of 0.07 s per voxel with either the 60× UPlanSApo or 100× UPlanFLN objectives. On average, we measured 3–10 cells of each strain from each cell culture. In case of environmental samples, we measured at least one cell. Standards of pure chemical substances were measured in water suspension. To prepare the matching references for biogenic crystals of uric acid, guanine monohydrate, xanthine, and their mixtures, the substances were dissolved in an aqueous solution (4 %) of dimethylamine (DMA) and dried on the quartz slide to allow recrystallization.

Data was analyzed using WITec Project FIVE Plus v5.1 software (WITec, Germany) to implement the following steps: cosmic ray removal, background subtraction, cropping of the spectral edges affected by detector margins, spectral unmixing with the true component analysis tool, and averaging of the mean spectrum, summarizing multiple measurements in order to optimize the signal-to-noise ratio for each single spectrum of the crystalline inclusions.

### Phylogenetic analyses

In an attempt to evaluate the role of nucleobase-cation symporter 1 (NCS1), nucleobase-ascorbate transporter (NAT), AzgA, and hypoxanthine-guanine phosphoribosyl transferase (HGPT) in purine crystal biocrystallization, we tested their phylogenetic distribution and the robustness of the phylogenetic placements using methods of molecular phylogenetics. We performed an extensive set of searches of eukaryotic and prokaryotic sequence databases. Using several representative sequences of each gene as queries, we performed a BLASTp search against 87 high-quality, well-annotated eukaryotic and prokaryotic genomes and transcriptomes. To exclude the possibility that absence of NCS1, NAT, and AzgA gene in predicted proteomes from Heterolobosea, Ciliophora, and Apicomplexa is caused by suboptimal protein prediction, we also checked the presence of their homologs (using TBLASTN search) in contigs from eight nucleotide genome assemblies representing the three groups. While HGPT homologs were easy to detect, TBLASTN did not detect any purine transporter genes. Thus, we can be confident that the absence of these genes in genomic data is not artificial. Datasets containing original sequences and their manually curated homologs, identified by BLASTp search, were included into initial datasets and aligned by MAFFT version 7. These alignments were used as inputs to build Hidden Markov Models for a final sensitive homolog search by HMMER3 software^26^ to identify candidate proteins from 742 eukaryotic genomes and transcriptomes included in the EukProt database of genome-scale predicted proteins across the diversity of eukaryotes^27^. To avoid bias introduced by contaminations or erroneous protein predictions in the EukProt database, we performed a preliminary set of Maximum-Likelihood phylogenetic analyses using IQ-TREE multicore version 1.6.10 ^28^ under LG4X model. Between every round, we manually inspected each tree to identify possible eukaryotic and prokaryotic contaminations. Suspicious sequences were used as queries against the NCBI database of non-redundant proteins (nr) and best blast hits were added to the dataset. Final gene datasets, free of contaminant sequences and in-paralogs (recent gene duplications that resulted in several homologs with almost identical sequence), were aligned by MAFFT ^29^. Alignments were manually edited in BioEdit ^30^, phylogenetic trees were constructed by the maximum likelihood method using RAxML ^31^, with LG+GAMMA+F model selected by Modelgenerator ^32^, and 200 nonparametric bootstrap analyses. Trimmed datasets of AzgA, NCS1, NCS2, NATs, and HGPRT protein families are stored on an online depository server, which will be accessible upon full article publication: https://figshare.com/s/ec36ff8263c1114d547a.

For assessing the distribution of equilibrative nucleoside transporter or solute carrier 29 (ENT, SLC29) and concentrative nucleoside transporter (CNT, SLC28), we used seed-sequences according to the references ^33^ as initial datasets aligned by MAFFT version 7. These alignments were used as inputs to build a profile HMM followed by an HMM search against 57 sequences of genomes using HMMER3 software ^26^.

## Data and materials availability

All data is available in the main text or the supplementary materials.

## Acknowledgments

We express our gratitude to Kateřina Schwartzerová, Jan Petrášek, Stanislav Vosolsobě, Lukáš Falteisek, Jana Krtková and Richard Dorrell for constructive discussions. English has been kindly corrected by William Bourland. Furthermore, we thank to Dovilė Barcytė, William Bourland, Antonio Calado, Dora Čertnerová, Yana Eglit, Ivan Fiala, Martina Hálová, Miroslav Hyliš, Dagmar Jirsová, Petr Kaštánek, Viktorie Kolátková, Alena Kubátová, Alexander Kudryavtsev, Frederik Leliaert, Julius Lukeš, Jan Mach, Joost Mansour, Jan Mourek, Yvonne Němcová, Fabrice Not, Vladimír Scholtz, Alastair Simpson, Pavel Škaloud, Jan Šťastný, Róbert Šuťák, Daria Tashyreva, Dana Savická, Jan Šobotník, Zdeněk Verner, Jan Votýpka for kindly providing cultures and taxonomic identifications. **Funding:** Financial support from the Czech Science Foundation (grants 17-06264S, 19-19297S, 20-16549Y, 21-03224S, and 21-26115S); Grant Agency of Charles University (grant 796217), Charles University Research Centre program No. 204069, European Regional Development Fund and the state budget of the Czech Republic, projects no. CZ.1.05/4.1.00/16.0340, CZ.1.05/4.1.00/16.0347, CZ.2.16/3.1.00/21515 and CZ.02.1.01/16_019/0000759, LM2018129.

## Author contributions

JP conceived the study, handled the cell cultures, performed the Raman measurements and data processing, prepared the graphics and videos and wrote the manuscript; TP and MO performed phylogenetic analyses and profiling; PM conceived the study, processed the data and corrected the manuscript; IČ provided the cell cultures and corrected the manuscript. All authors discussed and approved the manuscript.

## Competing interest declaration

Authors declare no competing interests.

**Supplementary Information** is available for this paper.

## Supplementary Text

### Results

Here we provide a detailed description of the results presented in Fig. 1 and Table S1. We examined representatives of all currently recognized eukaryotic supergroups ^34, 35^ for the presence and composition of purine inclusions, *i.e.,* guanine anhydride, guanine monohydrate, uric acid and xanthine. To the best of our knowledge, this is the first report on the occurrence of pure crystalline guanine monohydrate in any microorganism. Apart from purine inclusions being formed by pure substances (Fig. S1), we also found four species forming mixed crystals consisting of various proportions of guanine monohydrate, uric acid and/or xanthine (Fig. S2).

Guanine inclusions (Movie S1) were ubiquitous within the **SAR** clade (Stramenopiles, Alveolata, Rhizaria). In **Alveolata,** we confirmed, for the first time, that the morphologically prominent “polar granules” in unsporulated oocysts of the parasitic apicomplexan *Eimeria maxima* consist of guanine. Furthermore, we proved that inclusions in all observed alveolates, including parasitic apicomplexans (*Eimeria maxima*, *Psychodiella sergenti*), photoparasitic chromerids ^36^, and very diversified and ecologically important dinoflagellates and ciliates consist exclusively of guanine. Among dinoflagellates, our sampling included clades with species having plastids derived from diatoms (*Glenodinium foliaceum*) and also those having complex plastids of rhodophyte origin ^37^. Moreover, we also included the bloom-or red-tide-causing species *Heterocapsa triquetra* ^38^ and a fresh isolate of Symbiodiniaceae from the soft coral *Capnella imbricata*.

Compared to Alveolata, in **Stramonopiles** the situation is more complex. Guanine crystals predominate in most species, such as predatory actinophryids, bicosoecids, phototrophic and heterotrophic chrysophytes (*Synura hibernica*, *Spumella* sp., respectively), biotechnologically promising eustigmatophytes (*Nannochloropsis oculata, Eustigmatos* cf. *polyphem*) ^39, 40^, labyrinthulomycetes (*Schizochytrium* sp.) ^41^, raphidophytes (*Gonyostomum* sp.), and xanthophytes (*Botrydiopsis intercedens*, *Tribonema aequale*, *Xanthonema* sp.). In contrast, diatoms have a lower prevalence of crystalline inclusions than other stramenopiles and exhibit production of uric acid crystals, for example freshwater species of *Encyonema*, *Fragilaria*, *Navicula*, and *Pleurosigma*. Guanine crystals were detected in only a single marine/brackish diatom species (Naviculaceae gen. sp., *Seminavis-*like). Diatoms are known for a very complex nitrogen metabolism employing a urea cycle comparable to the one in animals ^42^. Crystalline inclusions were not detected in the parasitic *Blastocystis* (Opalinata) or in Oomycota. Guanine crystals predominated in the sampled **Rhizaria** (*i.e.,* Cercozoa). In Gromiida and in Retaria (Foraminifera, Acantharea, Polycystinea) we suspected the previously described structures called “stercomata” ^43^ to be crystals of purines, calcite or celestite (strontium sulfate). For now, we did not prove any of those with certainty due to low sampling of only three species of formalin-fixed cells of Acantharea and Foraminifera.

The more intricate crystals of **Haptista** may contain a mixture of both purines (Movie S2). Among haptophytes, *Emiliania huxleyi* ^44^, a model organism with ecological importance, possesses guanine crystals, whereas the biotechnologically important *Isochrysis* sp. ^41^ has xanthine crystals. In both cases, organisms were cultured in guanine-supplemented medium (Table 1). Centroplasthelida from freshwater habitats (*Rhaphidiophrys* sp.) possessed numerous guanine crystals. **Telonemida** (*Telonema* sp.) contained guanine inclusions when cultured in guanine-supplemented medium. Within **Cryptista (***Chroomonas* sp., *Cryptomonas* sp.) we showed that the highly refractile and taxonomically important Maupas body ^45^ consists of uric acid.

In **Archaeplastida** (Movie S3), guanine crystals were present in glaucophytes but were not detected in any of the nine sampled rhodophyte species. Guanine inclusions are consistently distributed throughout the UTC clade (Ulvophyceae, Chlorophyceae, and Trebouxiophyceae) except for the biotechnologically significant species, *Chlorella vulgaris* ^41^, which possess xanthine, and *Dictyosphaerium* sp. ^46^, which contains uric acid crystals. The arctic species *Chloromonas arctica* ^47^ possesses guanine crystals. Interestingly, *Chlamydomonas* in culture or environmental samples possesses guanine crystals in different life stages from flagellate to palmelloid. Xanthine is also present in the crystals of chlorodendrophytes, including another biotechnologically exploited species, *Tetraselmis subcordiformis* ^41^, whereas other marine chlorophyte counterparts contain guanine crystals (*e.g. Nephroselmis* sp.). The smallest free-living eukaryote, with the size of bacteria, *Ostreococcus tauri* (Mamiellophyceae) ^48^, did not show any crystalline inclusions even though they may contain starch as a storage polysaccharide. It is also possible that our methods had insufficient resolution to detect crystalline inclusions in these tiny cells. In stark contrast to Chlorophyta, the Streptophyta ^49^, notably including land plants (Embryophyta) together with the related microalgae, Zygnematophyceae and Klebsormidiophyceae, contain uric acid inclusions. In the case of *Mesotaenium caldariorum*, the predominant uric acid include admixtures of crystalline guanine monohydrate. However, other examined streptophytes such as *Chlorokybus atmophyticus* (Chlorokybophyceae) form guanine crystals. We detected no crystalline inclusions in *Mesostigma viride* (Mesostigmatophyceae). In Embryophyta, Coleochaetophyceae, and Zygnematophyceae we observed calcium oxalate inclusions instead of purine crystals (see details below).

Crystalline inclusions of varied chemical composition, primarily guanine and xanthine, occur commonly in **Amoebozoa** (Movie S4). Xanthine crystals, uncommon in other eukaryotes, are present in freshwater and marine *Mayorella* sp., in the facultatively parasitic model organism, *Acanthamoeba castellanii*, in the anaerobic model species *Mastigamoeba balamuthi*, and a model terrestrial slime mold, *Physarum polycephalum* ^50, 51^. By contrast, guanine was present in a fresh isolate of another slime mold, *Fuligo septica*, a different species of Archamoebae (*Mastigella eilhardi*), and in species of the common genera *Thecamoeba*, *Vannella*, and *Difflugia*.

We showed by Raman microscopy that, in **Opisthokonta,** uric acid, rather than guanine, is a common excretory product of nitrogen metabolism (*e.g.* in nematodes). Interestingly, we noticed guanine microcrystals inside swarming acoelomate gastrotrichs (Platyzoa) in a sample from a peat bog (Movie S5). Using Raman microscopy, we also confirmed that guanine crystals serve as a refractile layer on fish scales ^52^. In Fungi (Holomycota) we found crystalline guanine in *Candida albicans*, which has been tested for uptake of different purine compounds previously ^53^. However, we did not find any crystalline inclusions in *Saccharomyces cerevisiae*. We observed no crystalline inclusions in either intracellular parasites belonging to Microsporidia or in the free-living halophilic Choanoflagellata. We did not detect crystalline inclusions in the **Breviatea**, close relatives of opisthokonts.

In **Excavata** (Movie S6), the crystalline inclusions, present in all studied lineages, were exclusively composed of guanine, including: both heterotrophic (*Entosiphon* sp., *Rhabdomonas* sp.) and photosynthetic euglenids, freshwater and marine euglenids (*Euglena* sp.*, Eutreptiella gymnastica,* respectively), in free-living kinetoplastids (but not in the parasites, such as trypanosomatids), in deep-sea diplonemids (*Namystinia karyoxenos, Rhynchopus* sp.) ^54, 55^, aerotolerant heteroloboseans (*Naegleria gruberi*) ^56^, and strictly anaerobic metamonad symbionts of termites (*e.g. Macrotrichomonoides* sp. isolated from *Neotermes cubanus*) ^57^. In the diplonemid, *Flectonema* sp., guanine monohydrate was detected in the form of pure crystals, probably monocrystals, as their Raman spectra exhibit similar variability of relative intensities when excited by polarized light to that of synthetically prepared monocrystals.

We found guanine crystals in *Gefionella okellyi*, from the phylogenetically distinct clade of **Malawimonadida** ^58^. Among the **CRuMs** group (Collodictyonida, Rigifilida, and Mantamonadida) ^59^, we tested mantamonads but found no crystalline inclusions inside their cells, even after exposing them to guanine-enriched culture medium. Lastly, crystalline guanine was found in *Hemimastix kukwesjijk*, the only representative sampled from **Hemimastigophora** ^58^, a separate, deep-branching lineage, recently described as a new eukaryotic supergroup (Movie S7).

In addition to the predominant purine crystalline inclusions (80 %), there are also other types. Surprisingly, calcite (calcium carbonate – CaCO3) was present in diatoms ^60^, in which amorphous silica (silicon dioxide – SiO2), that does not polarize light, is the main component of their biomineralized frustules^61^. Calcification (Fig. S3, Tab. S1) occurred in the seaweeds tested, including rhodophytes, ulvophytes and phaeophytes, indicating that this process occurs very commonly in marine environments ^62^. Similarly, calcified shells of foramiferans and the calcite scales on the cell surface of the haptophytic coccolithophore *Emiliania huxleyi* also strongly polarized light ^63^. Massive calcite incrustations occurred on surface of filamentous algae, *e.g. Oedogonium* sp., that has been previously described as crystal jewels in other freshwater filamentous algae ^64^. We confirmed strontianite (strontium carbonate – SrCO3) in the green alga *Tetraselmis*, previously reported elsewhere using different methodology ^65^.

Crystalline sulfates occurred in various species but only rarely (Fig. S4). Similarly, in the light-polarizing armor of Acantharea, we confirmed the presence of celestite (strontium sulfate – SrSO_4_) ^66, 67^. Interestingly, we found baryte (barium sulfate – BaSO_4_) in three species of zygnematophytes (*Closterium peracerosum-strigosum-littorale* complex, *Cosmarium* sp., and *Spirogyra* sp.), for two of which it has been previously reported ^68, 69^. In laboratory cultures and environmental isolates of *Saccamoeba* sp. we found an unprecedently complex spectrum corresponding to numerous crystalline inclusions lacking light-polarization features. The dominating peak at 998 cm^−1^ (Fig. S4, Tab. S1) resembles either a signal of sulfates or aromatic compounds. The rest of the spectrum also shows lipid-like organic matter, thus, the crystals may be formed by a complex mixture of lipids and sulfates.

Calcium oxalate monohydrate (CaC_2_O_4_ · H_2_O) crystals were restricted to closely related streptophytic algae and land plants: Coleochaetophyceae, Zygnematophyceae (*Cylindrocystis* sp.), and Embryophyta (*e.g.* plant models *Physcomitrella patens*, *Nicotiana tabacum*) together with calcium oxalate dihydrate (CaC_2_O_4_ · 2H_2_O) commonly found in Embryophyta (Fig. S5, Tab. S1). In the Embryophyta, deposition of calcium oxalate is already known to occur under stress conditions ^70^.

Unexpected light-polarizing lipophilic inclusions unreported until now, occurred in some of examined samples. Compared to carotene crystals observed in the model plant *Arabidopsis thaliana* ^71^, we found possibly similar structures in aerophytic ulvophytes (*Trentepohlia* sp., *Scotinosphaera gibberosa*) and freshwater cyanobacteria (*Oscillatoria* sp.), comprising a mixture of carotenoids and lipids containing sterols and fatty acids (Fig. S6, Table S1, Movie S8). Similarly, in the green parasitic alga, *Phyllosiphon arisari* ^72^ and a symbiont of lichens, *Symbiochloris tschermakiae* ^73^, their light-polarizing lipophilic crystals were a complex mixture of lipids, with a high proportion of unsaturated fatty acids in the former, and saturated fatty acids in the latter. Faintly light-polarizing lipophilic structures in amoebozoans (*Entamoeba histolytica*) resembled those mentioned above or contained a surplus of sterol compounds (*Acanthamoeba castellanii* and *Mastigamoeba balamuthi*). Some of them may be of a crystalline nature but this requires further evidence. All lipophilic light-polarizing structures tended to melt under prolonged illumination by a focused laser beam (ca 20 mW power) during the Raman measurements.

Refractile structures such as storage polysaccharides, *i.e.,* starch or chrysolaminarin ^74, 75^, or aerotopes, air-filled vesicles with a reflective interface inside the cells of cyanobacteria ^76^, might be confused with birefringent crystals. High-intensity light-polarization also occurs in the thick cellulose cell walls of ulvophytes, zygnematophytes, rhodophytes, glaucophytes and others, a phenomenon best-studied in plants ^77^, or in starch, a storage polysaccharide of Archaeplastida and in chrysolaminarin, storage polysaccharide of SAR (Fig. S7, Table S1).

### Comments on Raman spectra analysis

In some protists, a monohydrate of guanine in crystalline form was detected (Figs. S1 and S2). To the best of our knowledge, this is the first report confirming the occurrence of crystalline guanine monohydrate in any microorganism. To date, only a β-polymorph of crystalline guanine anhydride was detected in various organisms, including some microalgae and protists ^4, 5, 8^. In *Flectonema* sp., guanine monohydrate was detected in the form of pure crystals (Fig. S1), probably monocrystals. We observed a few inclusions formed by purine mixtures, which has also been reported from *Paramecium* ^4^. In the case of *Mesotaenium caldariorum*, crystalline guanine monohydrate seems to be only a minor admixture in the more abundant uric acid (Fig. S2), however its presence was demonstrated by spectral similarity with synthetically prepared samples containing both uric acid and guanine monohydrate. The inverse proportion of the two compounds has been found in *Tetraselmis subcordiformis*. *Isochrysis* sp. and *Chroomonas* sp. exhibited crystalline mixtures of xanthine and guanine monohydrate.

### Comments on phylogenetic analyses

**The nucleobase cation symporter-1 (NCS1) family** (Fig. S10) of secondary active transport proteins includes proteins from prokaryotes and several lineages of eukaryotes. We recovered NCS1 eukaryotic paralogs – fungal Fcy type, fungal Fur type, algal type, and plant type as previously described ^78^. Besides, we also identified four as-yet-unknown eukaryotic paralogs that we marked as NCS1 A–D. Distribution of NCS1 in eukaryotes is extremely patchy as summarized on Fig. S8. We conclude that NCS1 transporter has been acquired by eukaryotes several times independently. Fungal Fcy type is present in various Fungi (Ascomycetes, Basidiomycetes, Gonapodya). Some Fungi even contain several distant paralogs of this gene (e.g. *Aspergillus* and *Candida*). Interestingly, homologs from Oomycota form a clade within fungal sequences, branching sister to *Aspergillus* sequences with full statistical support. Thus, our analysis strongly indicates lateral transfer of this gene from Fungi (supergroup Obazoa) to Oomycota (supergroup SAR). We identified fungal Fcy type of NCS1 transporter also in Ktedonobacteria (Chloroflexi) and two groups of Proteobacteria (Betaproteobacteria and Gammaproteobacteria). Bacterial homologs of fungal Fcy form a clade branching within fungal sequences but, in this case, with relatively low bootstrap support. Besides, the whole clade of fungal Fcy sequences shows strong affinity to eubacterial permeases including those from Gammaproteobacteria and Betaproteobacteria, so in this case, lateral transfer of Fcy from Fungi to Eubacteria is less convincing. We also identified an ecological relationship between eukaryotes that possess fungal Fcy gene, all of which are adapted to extract nutrients from plants. The Fur type of NCS1 is present exclusively in Basidiomycetes and Ascomycetes (Fungi). The source organism for Fur type is unclear. The algal type of NCS1 is present in *Nanochloropsis* (SAR), Rhodophyta, and Chlorophyta (Archaeplastida). It also has an uncertain origin. Plant type NCS1 was detected in Chlorophyta and Streptophyta (closely related lineages of the supergroup Archaeplastida). Surprisingly, this gene is also present in Rhodelphidia, another deep-branching lineage of the supergroup Archaeplastida. The gene has unclear origin, although it shows affinity to Proteobacteria with low statistical support.

We also identified four novel clades of eukaryotic NCS1 transporters. One of them is present only in dinoflagellates and shows a close relationship to cytosine permease from Actinobacteria (with full bootstrap support); another is in *Chromera*, diatoms, and dinoflagellates; it shows affinity to Bacteroidetes and Planctomycetes with full bootstrap support. The other clade contains homologs from choanoflagellates, Hemimastigophora, telonemids, dinoflagellates, Chlorophyta, and malawimonadids. The phylogenetic position of *Incisomonas marina* (Stramenopiles) within choanoflagellates is probably due to contamination, also most likely in the case of *Paulinella* (Rhizaria) within Chlorophyta. Finally, two sequences from unrelated amoebozoans (*Filamoeba* and *Vermamoeba*) form a robust clade with no affinity to any other group, forming another eukaryotic NCS1 clade.

The **AzgA** gene encodes a **hypoxanthine-adenine-guanine transporter** (Fig. S9) that is present in all main groups of eukaryotes except supergroup Amoebozoa, and it is also missing in metazoans. Our analysis convincingly shows that eukaryotic AzgA has a single origin in eukaryotes and has been probably present in two paralogs in the last eukaryotic common ancestor. We named those paralogs AzgA A and B. Besides, the analysis indicates a series of gene duplications during evolution of certain groups as seen in Chlorophyta and Dinoflagellata.

**Nucleobase-Ascorbate Transporter (NAT) protein family** (Fig. S11) is an extensively studied group of proteins. All bacterial NATs are H^+^ symporters highly specific for either uracil or xanthine or uric acid. The fungal and plant members are H^+^ symporters specific either for xanthine-uric acid, or for adenine-guanine-hypoxanthine-uracil. In contrast to the microbial and plant proteins, most functionally characterized mammalian NATs are highly specific for L-ascorbate/Na^+^. However, the rSNBT1 NAT transporter from a rat is specific for nucleobases ^19^. Our analysis convincingly shows that NAT proteins were introduced to eukaryotes at least four times independently (NAT A–D) and have their closest homologs in eubacteria, NAT B and D with high statistical support. Metazoan and plant NATs both belong to the NAT A clade that is, together with NAT C, the most widespread NAT gene in eukaryotes. In contrast, NAT B is present only in Fungi and dictyostelids; NAD D is unique for *Tritrichomonas* and it was established by horizontal gene transfer from Firmicutes (it is encoded on tritrichomonas-like genomic contig, so it is not contamination). Interestingly, some well-supported eukaryotic clades are not congruent with eukaryotic phylogeny confounding interpretation of the descent of this protein family. In some cases, it might be explained by eukaryote-to-eukaryote horizontal gene transfers (*e.g.,* from fungi to dictyostelids in NAT B).

**Hypoxanthine-guanine phosphoribosyl transferase (HGPT)** (Fig. S12), omnipresent in eukaryotes, can also be found in Eubacteria, Archaea and, surprisingly enough, we detected HGPT homologs also in Nucleocytoviricota genomes. Because it is a relatively short and divergent protein, we were unable to resolve its detailed phylogeny. Based on previously introduced nomenclature ^79^, we distinguished a clade of “fungal HGPT” which is very divergent from the other “classical HGPTs”. However, we were able to find homologs of “fungal HGPT” in virtually all eukaryotic supergroups.

---

**Fig. S1.**
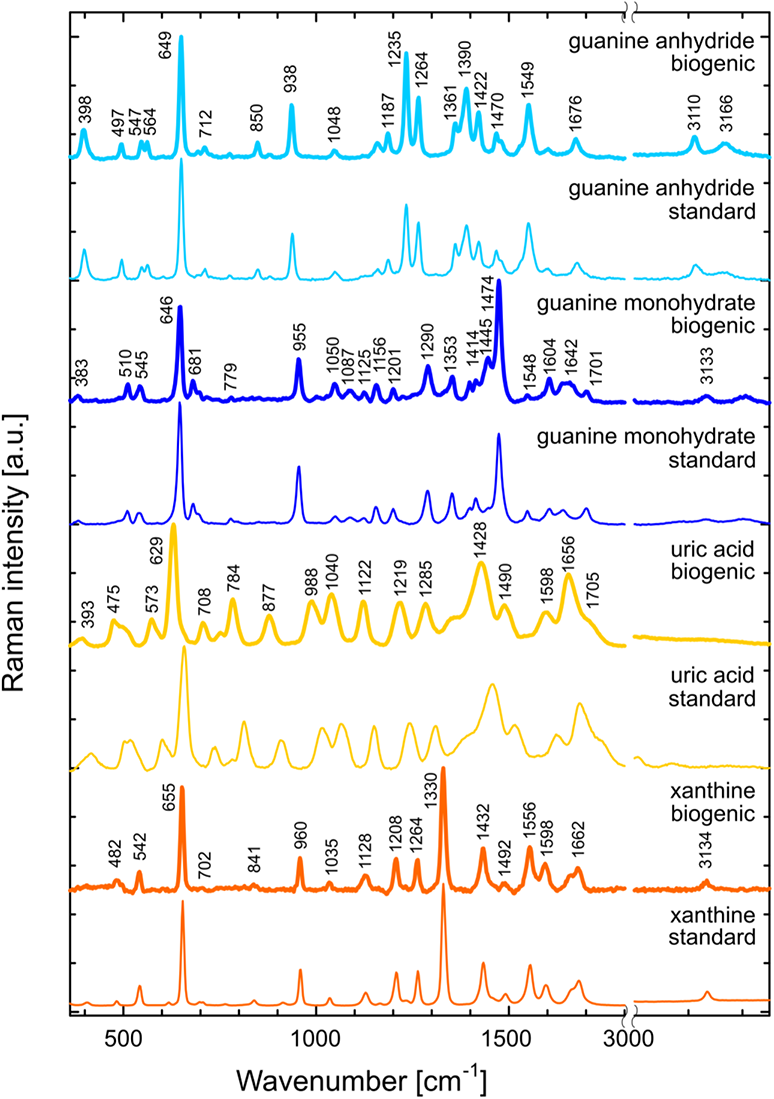
Representative Raman spectra of biogenic purine crystalline inclusions measured in biological species followed by their respective standards of pure chemical compounds. Biogenic crystals have been spectrally extracted directly from measured species. Standards of guanine anhydride and xanthine have been measured as suspension of pure compounds in water, guanine monohydrate and uric acid has been recrystallized from 4% dimethylamine water solution.

**Fig. S2.**
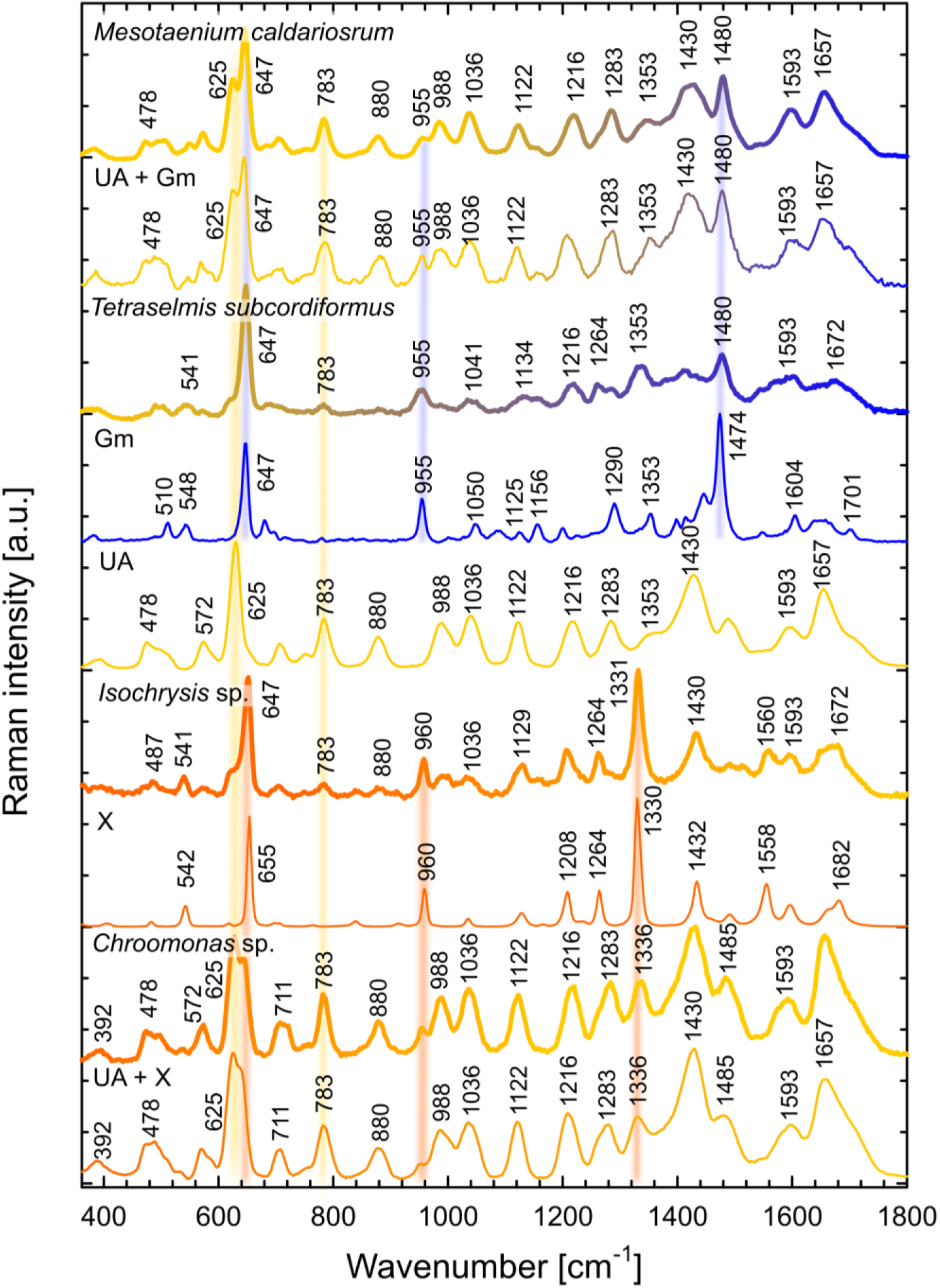
Representative Raman spectra of biogenic purine inclusions forming mixtures of uric acid (UA) and guanine monohydrate (Gm) in different proportions with a dominance of the former in the case of *Mesotaenium caldariorum*, or the latter in *Tetraselmis subcordiformis*. Xanthine (X) dominates the crystals with uric acid admixtures found in *Isochrysis* sp., whereas *Chroomonas* sp. has higher proportions of uric acid over xanthine. The reference spectra of pure substances are shown along with those of mixtures of “UA + Gm” and “UA + X” recrystallized from 4% dimethylamine solution, as Raman spectra of purine mixtures exhibit some spectral shifts and changes in relative intensities compared to the pure substances.

**Fig. S3.**
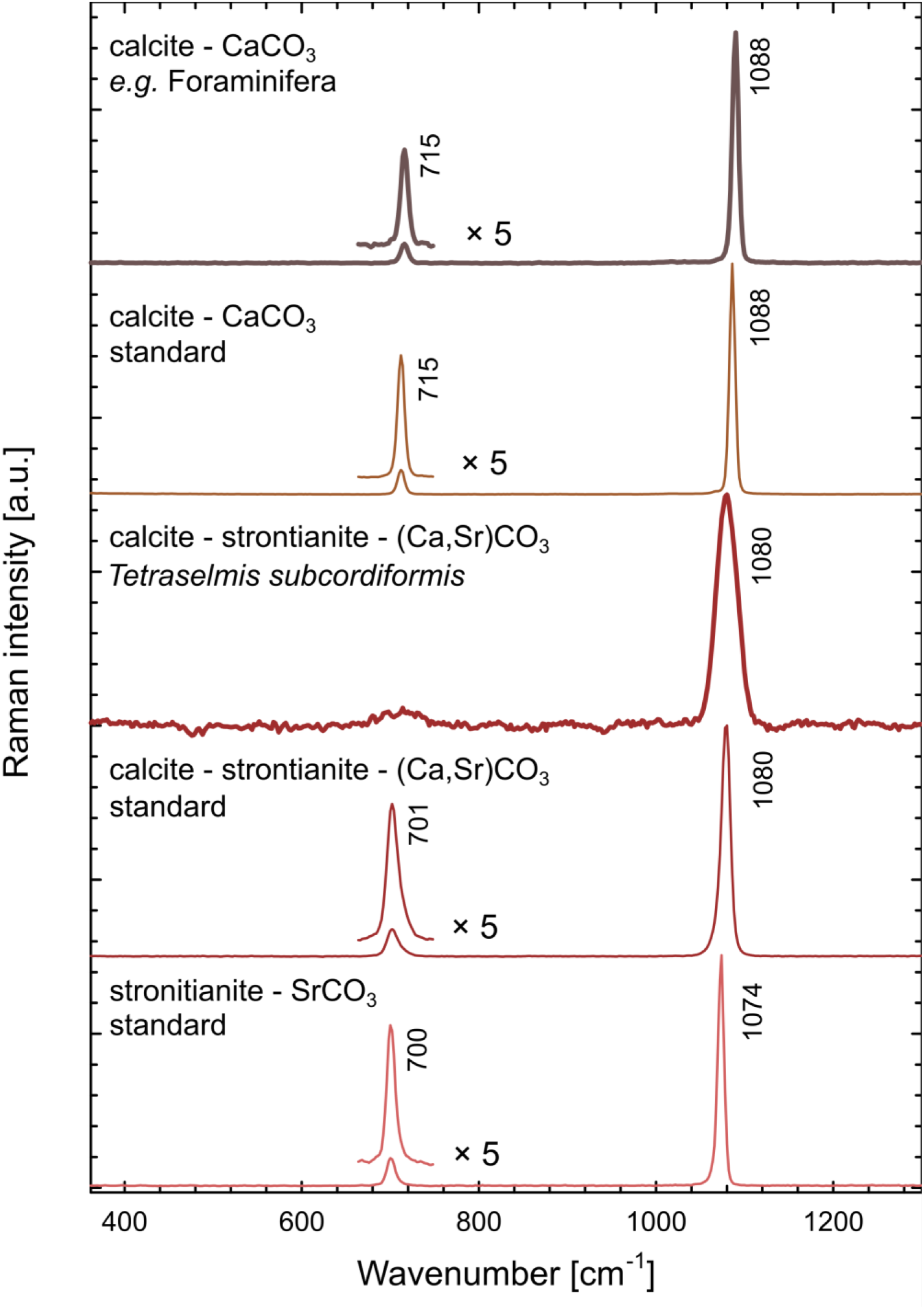
Representative Raman spectra of carbonate minerals observed in inspected species, all of them strongly polarize light: calcite or calcium carbonate (CaCO_3_) found in various diatoms, Foraminifera, multicellular Rhodophytes and seaweed with respective standard and calcite with admixtures of strontianite (SrCO_3_) or strontium carbonate (Ca,Sr)CO_3_ found in *Tetraselmis* sp. with respective standards of pure chemical substances.

**Fig. S4.**
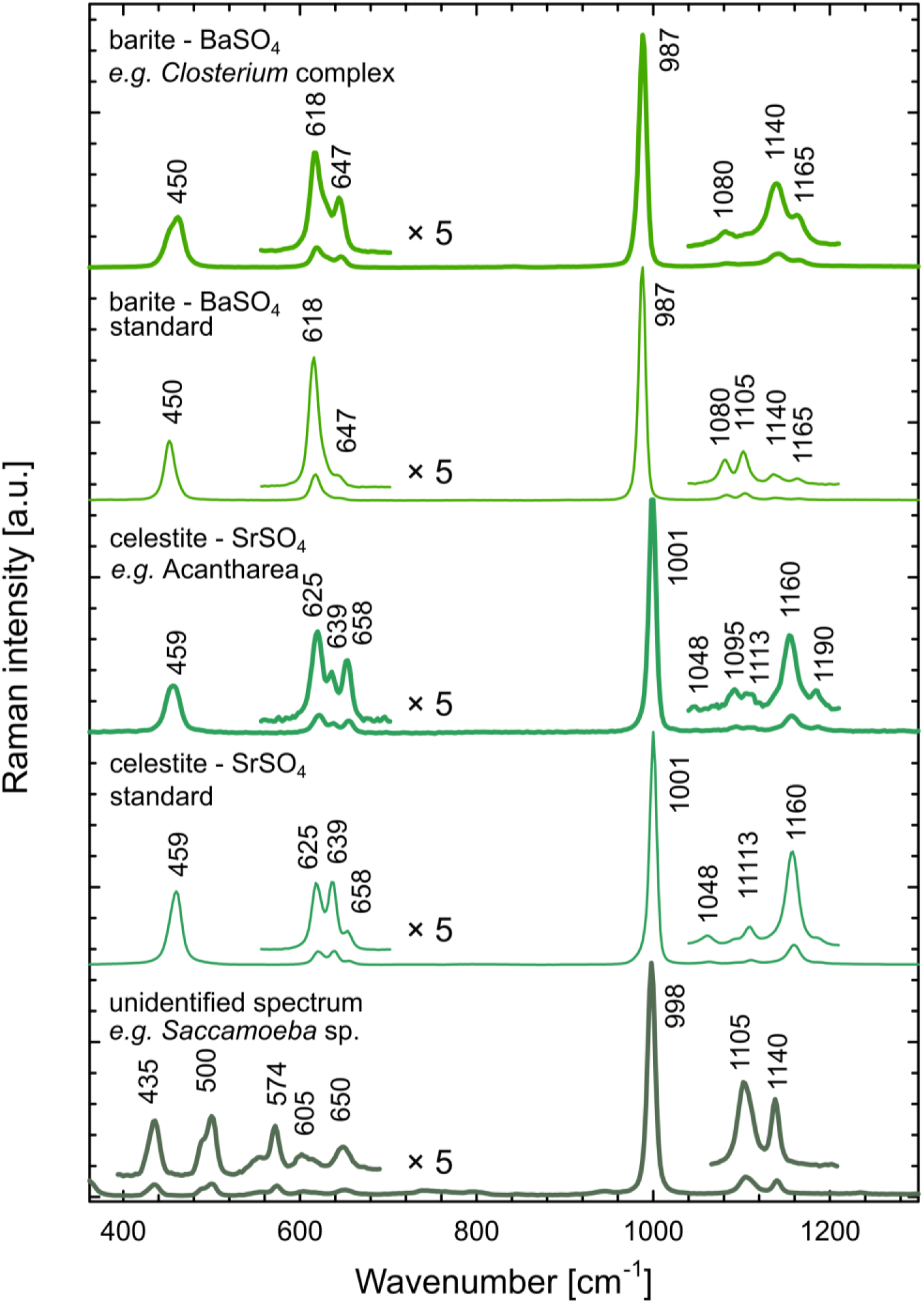
Representative Raman spectra of sulfate minerals observed in inspected species followed by respective standards of pure chemical substances, all of them faintly polarize light: baryte (BaSO_4_) found in *Closterium peracerosum-strigosum-littorale* complex, *Cosmarium* sp., *Spirogyra* sp., celestite (SrSO_4_) found in skeletons of Acantharea, and unidentified spectra of sulfate resembling minerals mixed with lipophilic organic matter found in *Saccamoeba* sp. Raman spectra of sulfates show variability in dependence of crystal orientation and minor admixtures of other salts.

**Fig. S5.**
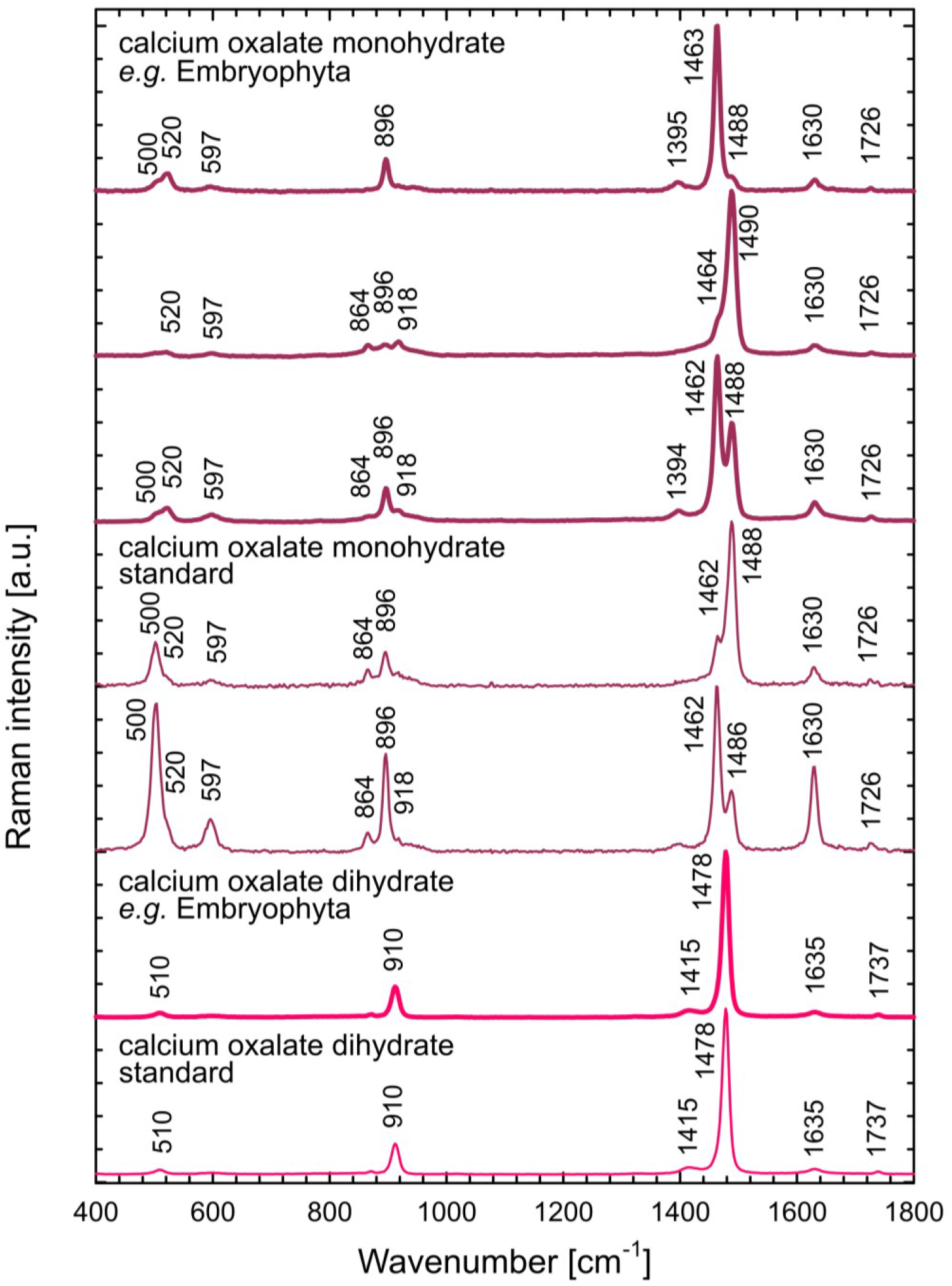
Representative Raman spectra of organic crystals polarizing light observed in inspected species followed by respective standards of pure chemical substances. The crystals of calcium oxalate monohydrate strongly polarize Raman signal – the major peaks interchange their relative intensities according to the crystal orientation with respect of polarization plane of the excitation beam. It was found in Coleochaetophyceae, Zygnematophyceae, Embryophyta. The calcium oxalate dihydrate was found in Embryophyta.

**Fig. S6.**
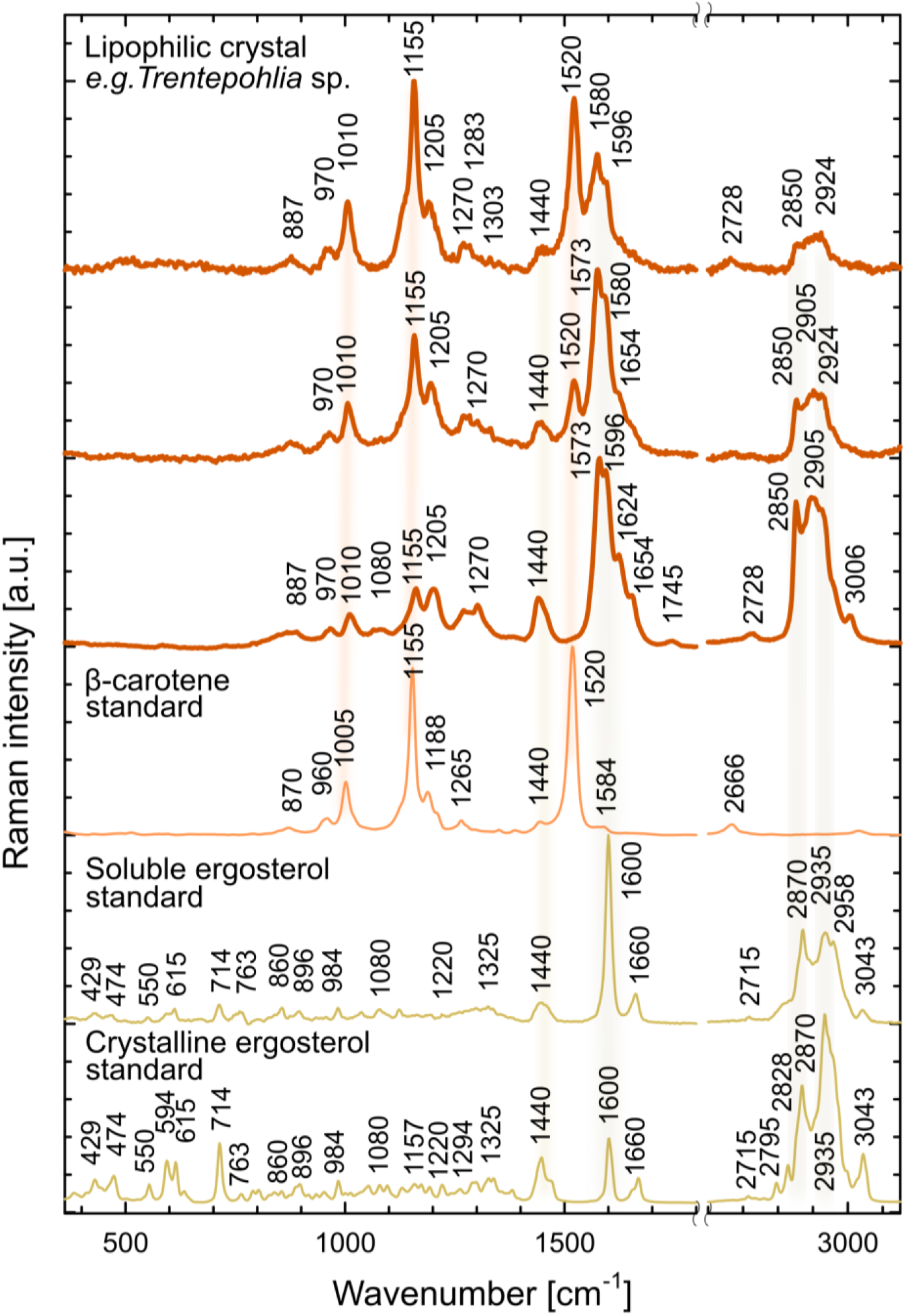
Representative Raman spectra of organic crystals polarizing light observed in inspected species followed by respective standards of pure chemical substances: lipophilic crystalline mixtures of carotenoids and sterols found in ulvophytes *Trentepohlia* sp. and *Scotinosphaera giberosa*, rhodophyte *Asterocytis ramonsa* and cyanobacteria *Oscillatoria* sp. Raman measurements using a high-intensity laser beam lead to photo-degradation of carotenoids present in the structure, thus their Raman signal decreases over time and allows observation of other admixtures in greater detail. We failed to find a precisely matching standard for sterols forming these lipophilic crystals; the biogenic crystals may contain a complex mixture of various chemical species.

**Fig. S7.**
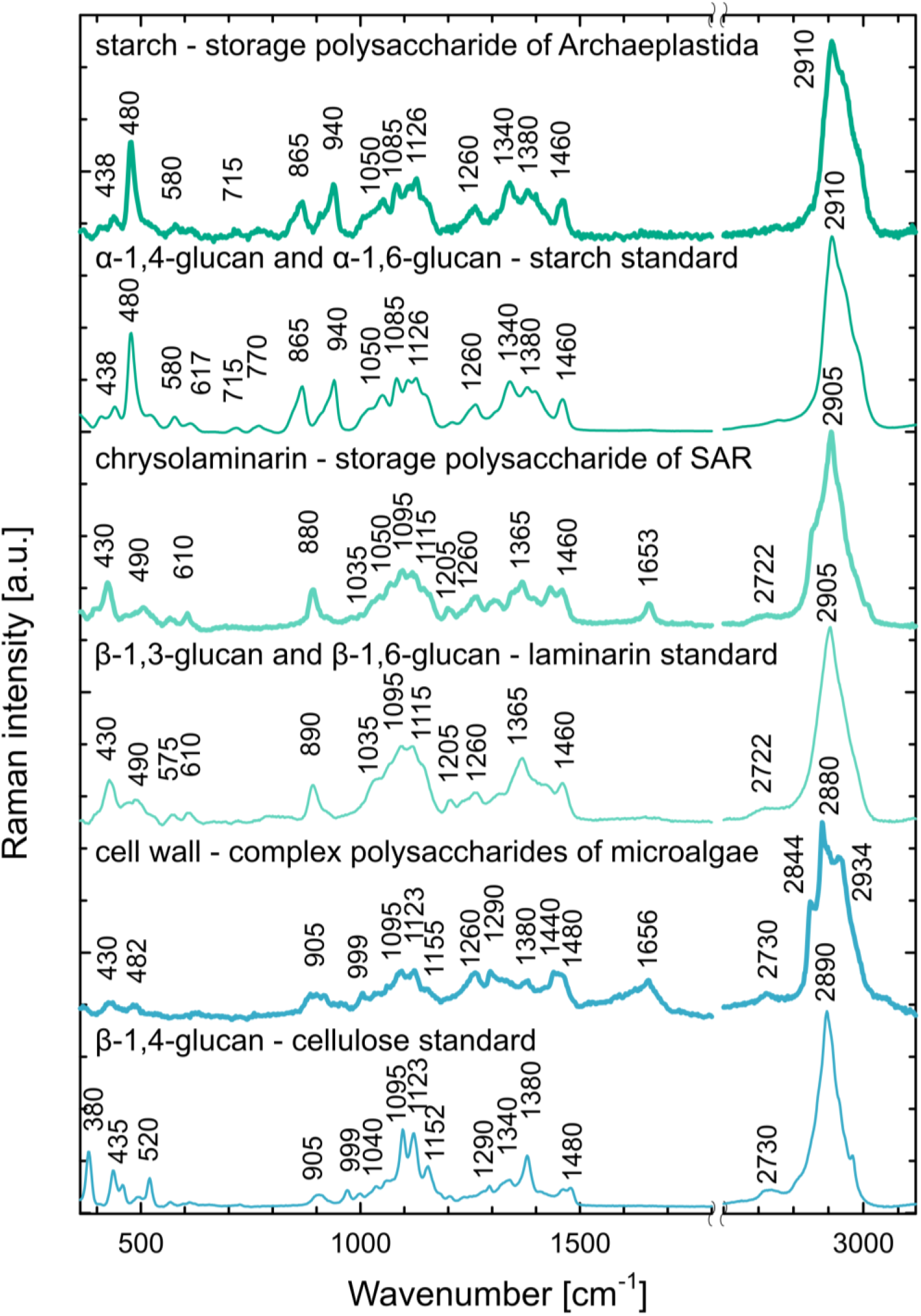
Representative Raman spectra of light polarizing polysaccharides in inspected species with respective standards of pure chemical substances: starch – storage polysaccharide of α-1,4-glucan and α-1,6-glucan found in Archaeplastida, chrysolaminarin – storage polysaccharide of β-1,3-glucan and β-1,6-glucan found in SAR, cellulose – structure polysaccharide of β-1,4-glucan forming cell walls of various microalgae (both Archaeplastida and SAR).

**Fig. S8.**
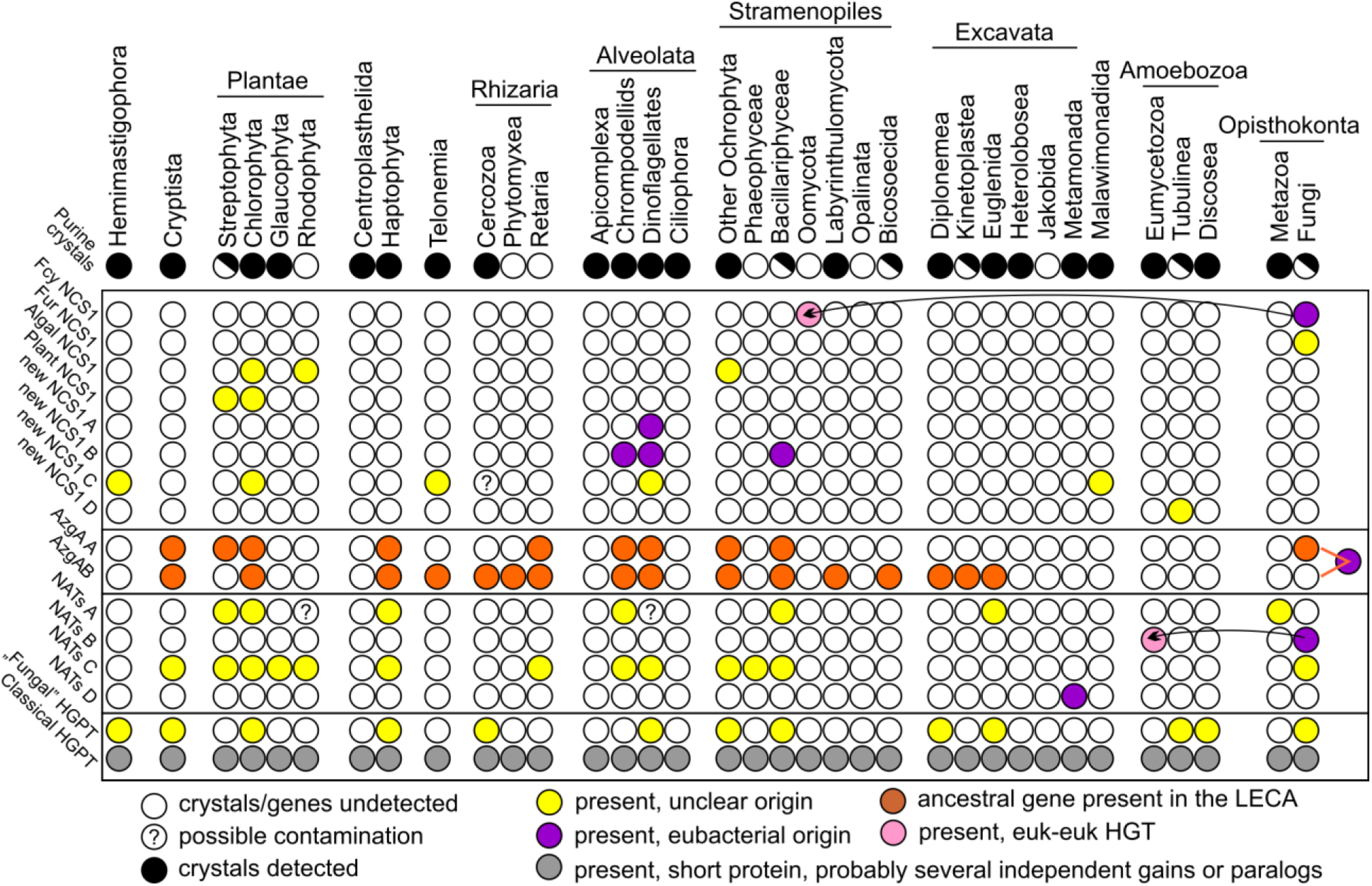
Summary table of phylogenetic distribution of the purine transporters: NCS1, NAT, AzgA, and the metabolic enzyme of salvage pathway – HGPT. There are notions on horizontal gene transfer (HGT) in two cases. In the case of AzgA we anticipate a possible origin in last eukaryotic common ancestor (LECA).

**Fig. S9.**
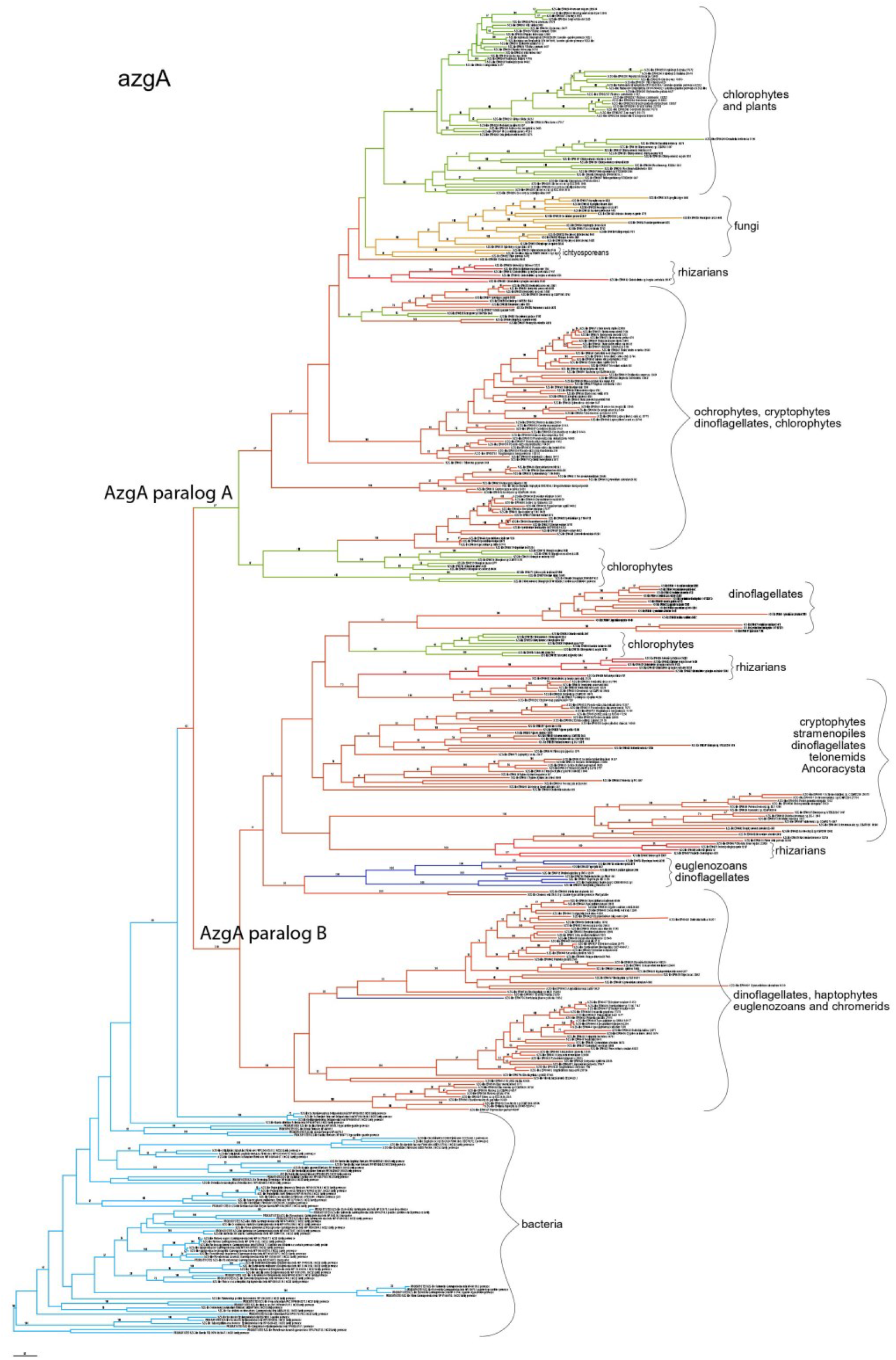
Maximum likelihood tree of AzgA as inferred from amino acid sequences (395 aa). Numbers above branches indicate ML bootstrap support (200 replicates).

**Fig. S10.**
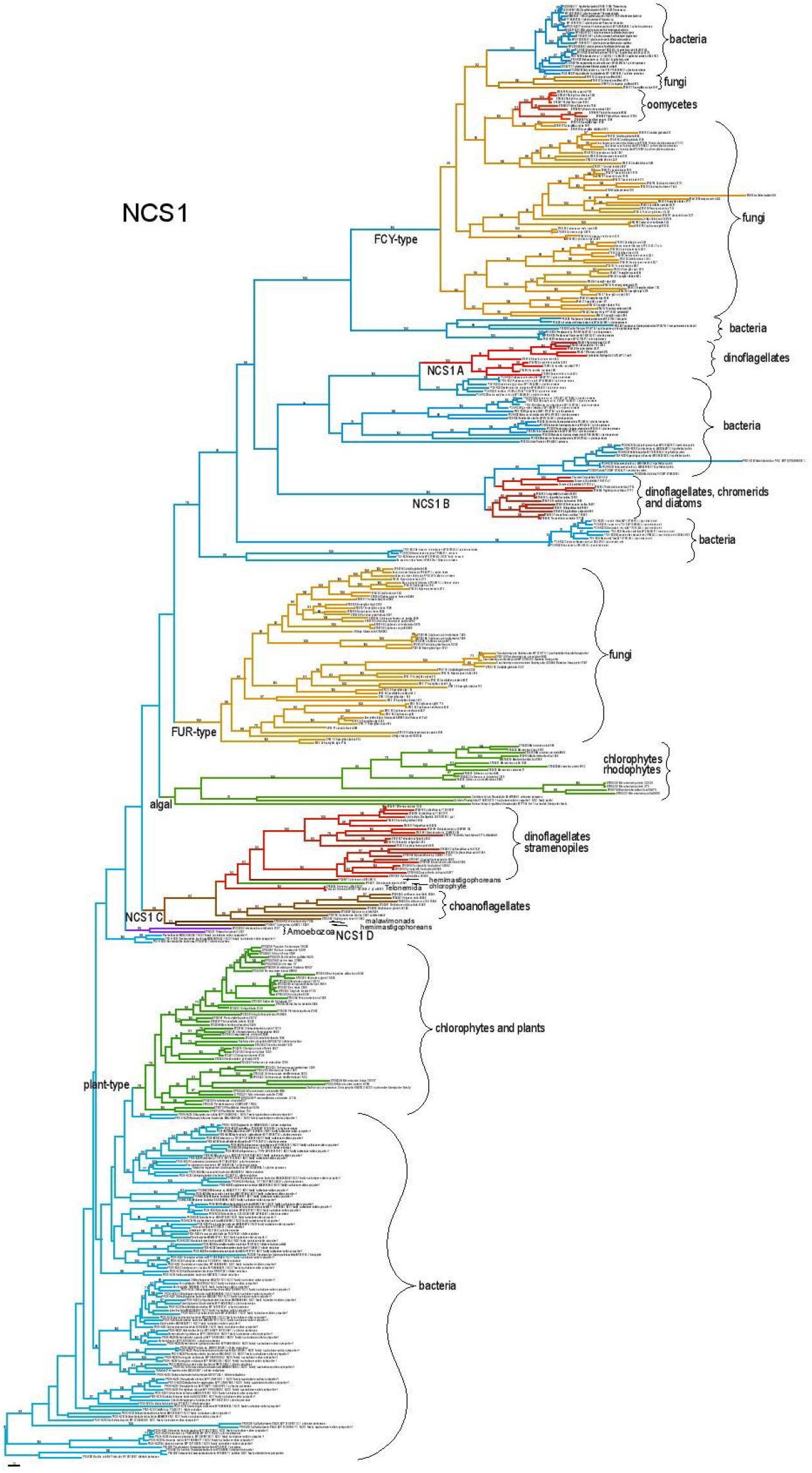
Maximum likelihood tree of nucleobase-cation symporter 1 (NCS1) as inferred from amino acid sequences (463 aa positions). Numbers above branches indicate bootstrap support (200 replicates).

**Fig. S11.**
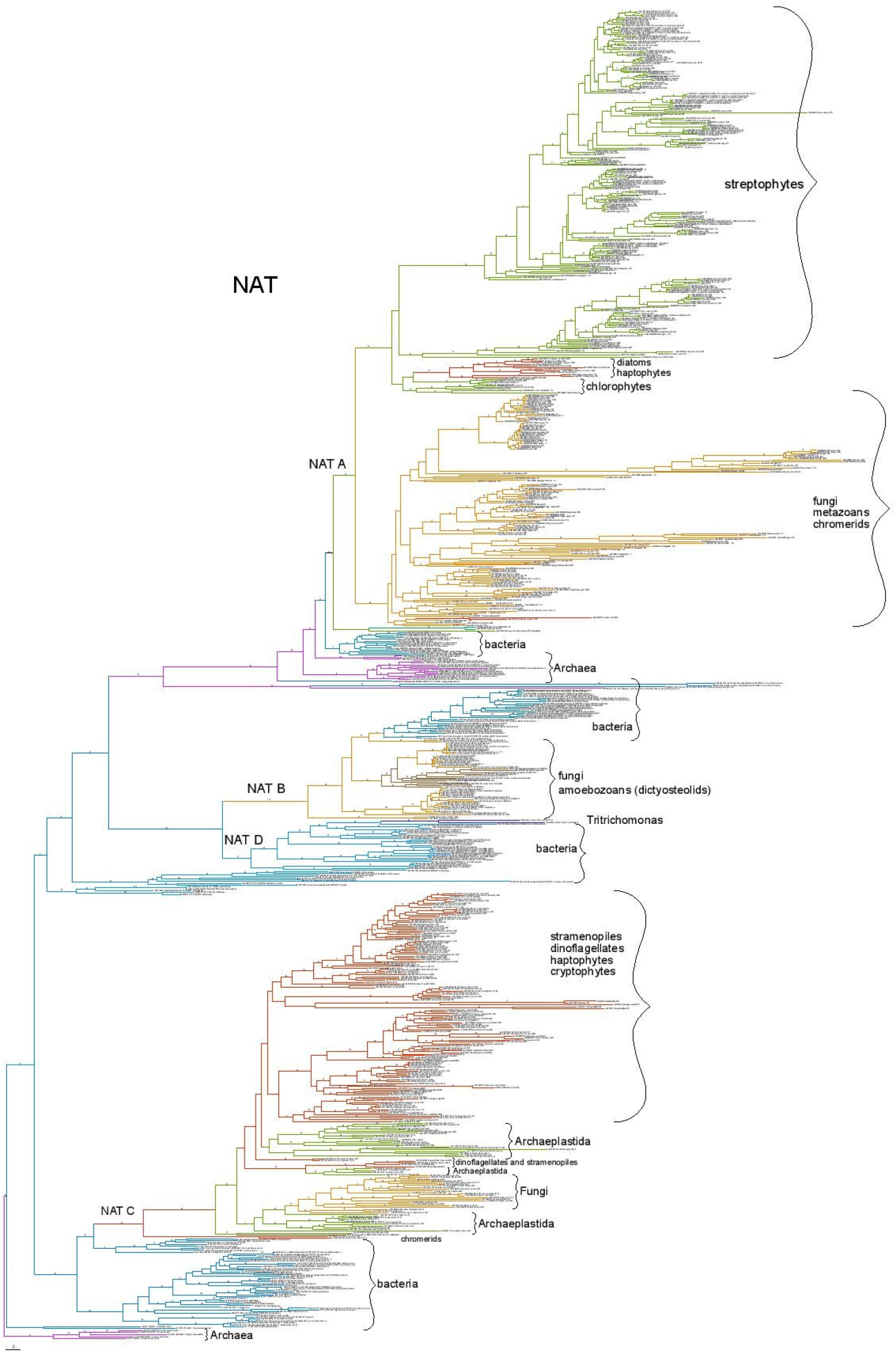
Maximum likelihood tree of nucleobase-ascorbate transporter (NAT) as inferred from amino acid sequences (385 aa). Numbers above branches indicate bootstrap support (200 replicates).

**Fig. S12.**
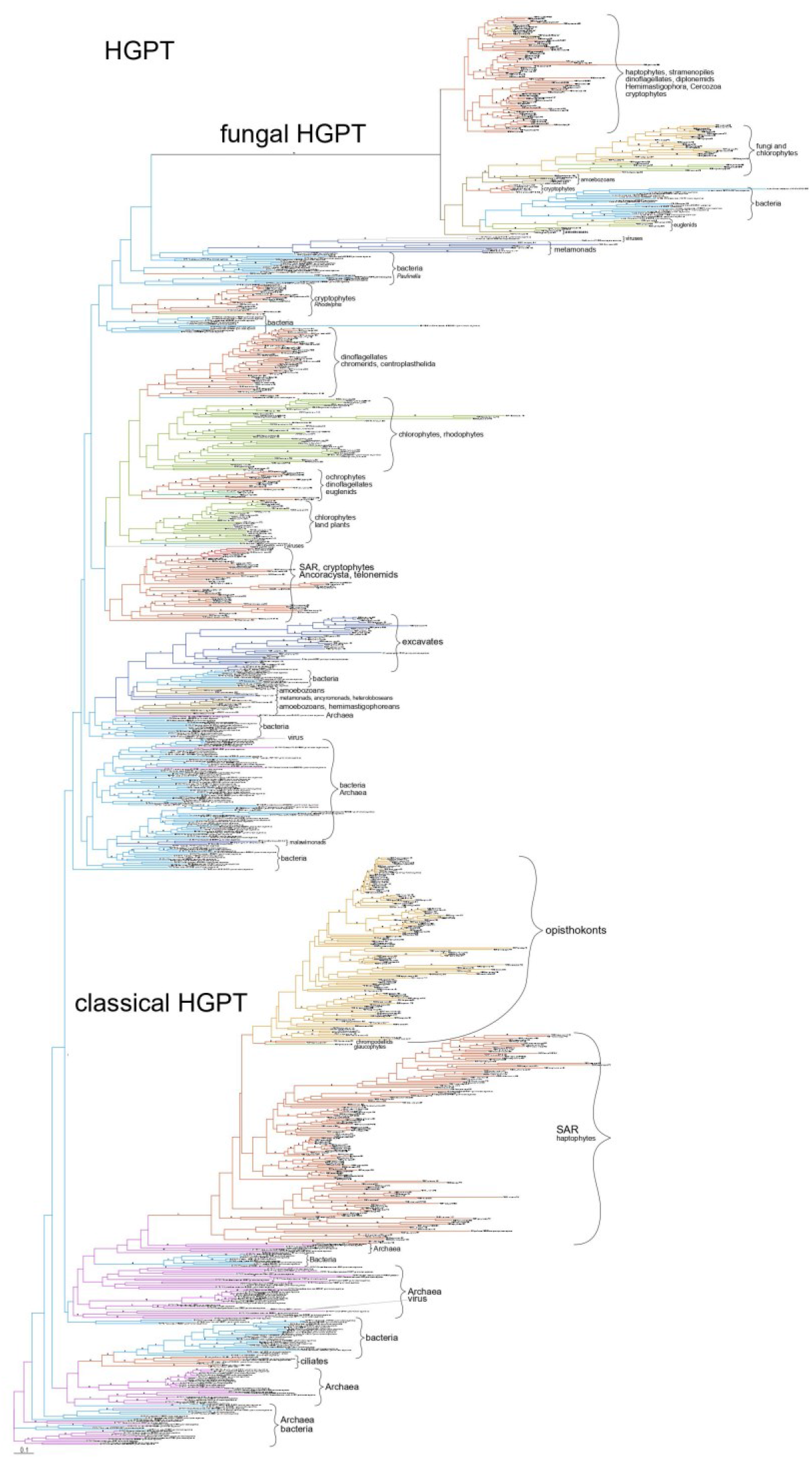
Maximum likelihood tree of hypoxanthine-guanine phosphoribosyl transferase (HGPT) as inferred from amino acid sequences (171 aa). Numbers above branches indicate bootstrap support (200 replicates).

**Table S1.** Overview of the cell inclusions present in eukaryotes and cyanobacteria and their identification *via* Raman microscopy and light polarization. Species are listed according to their taxonomic classification and then alphabetically – species positively tested for purine inclusions (in black) and not positively tested for purine inclusions (in grey); capital letters in brackets refer to the phylogeny scheme in Fig. 1; (b) stands for biotechnologically important species and (m) for model organism. **Habitat and-trophy:** E – Endobiont, F – freshwater, Hp – halophilic, M – marine, S – snow, T – terrestrial; a – autotroph, h – heterotroph, p – parasite. **Culture collections:** ATCC – American Type Culture Collection, Manassas, USA; CAUP – Culture Collection of Algae of Charles University in Prague, Czech Republic; CCALA – Culture collection of Autotrophic Organisms of the Institute of Botany of the AS CR, Třeboň, Czech Republic; CCAP – Culture Collection of Algae and Protozoa, Orban, UK; CCF – Culture collection of Fungi, Praha, Czech Republic; CCMP – Culture Collection of Marine Phytoplankton – part of NCMA: NCMA – National Center for Marine Algae and Microbiota, Bigelow, USA; CRC – Chlamydomonas Resource Center, University of Minnesota, St. Paul, USA; CVCC – China Veterinary Culture Collection Center, National Control Institute of Veterinary Bio-products and Pharmaceutical (NCIVBP), Ministry of Agriculture, Beijing, China; DBM – Department of Biochemistry and Microbiology at University of Chemistry and Technology, Prague, Czech Republic; EUROSCARF – EUROpean Saccharomyces Cerevisiae Archive for Functional analysis, Scientific Research and Development GmbH, Oberursel, Germany; NIES – National Institute for Environmental Studies, Tsukuba, Japan; NCMA – CCMP - National Center for Marine Algae and Microbiota, Culture Collection of Marine Phytoplankton, Bigelow, USA; priv – private collection; RCC – Roscoff Culture Collection, Roscoff, France; SAG – Culture Collection of Algae at Goettingen University, Germany; UTEX – University of Texas at Austin, USA **Cultivation method:** – – no cultivation prior to observations and measurements (in case of environmental samples) ASW-D – Artificial Sea Water for Diplonemids according to ^80^, 20 °C ASW-L – Artificial Sea Water for Labyrinthulomycetes: 3,5% artificial sea water supplied with 1 g of yeast extract and 10 g of glucose, 25 °C BBM – Basal Bold’s Medium according to ^81^, 20–25 °C, 16:8 h light:dark cycle BHI – Brain Heart Infusion medium as described in ^82^ DY-IV – medium according to ^83^, 15 °C, 12:12 h light:dark cycle f/2 – medium according to ^84^, 20–25 °C, 16:8 h light:dark cycle f/2B – f/2 medium according to ^84^ with addition of sterile barley grain, 20–25 °C Fw – ATCC 802 Sonneborn’s Paramecium medium, 20–25 °C FwAM – Fresh Water Amoeba Medium – ATCC medium 997, 20–25 °C FwB1 – 25% ATCC 802 Sonneborn’s Paramecium medium, with addition of sterile barley grain, 20–25 °C FwB2 – 25% ATCC 802 Sonneborn’s Paramecium medium, with addition of sterile barley grain, 17 °C HM-V – Halophiles Medium number V according to ^85^ with addition of sterile barley grain, 20– 25 °C JM – ATCC 1525, 25 °C LB – 3% Lysogeny Broth medium according to ^86^, 25 °C LBM – 3% Lysogeny Broth medium according to ^86^ in Artificial Sea Water, 25 °C M7 – medium according to ^87^, 25 °C MAM – Marine Amoeba Medium – ATCC medium 994, 25 °C MEA – Malt Extract Agar according to ^88^, 25 °C MS-1 – Murashige and Skoog liquid medium according to ^89^, 25 °C, 16:8 h light:dark cycle MS-2 – Murashige and Skoog solid medium according to ^89^, 25 °C, 16:8 h light:dark cycle Neff – medium of Neff and Neff according to ^90^, 25 °C OG – oat grain on cellulose paper kept in 100% humidity, 25 °C PDA – Potato Dextrose Agar according to ^91^, 25 °C PYGC – Proteose peptone – Yeast extract – Glucose–Cysteine medium according to ^92^, 25 °C TYI – ATCC 2695 Keister’s Modified TYI-S-33 medium, 37 °C TYM – medium modified according to ^93^ by an overlay of inactivated horse serum, 25 °C YPAD – ATCC 1069 medium, 37 °C † – fixed cells * – supplementary guanine added (approximately 30 µM final concentration), sterilized by 0.22µm filtration ^#^ – same species has been measured previously for publication in ^8^

– same species has been measured previously for publication in <sup>8</sup>
| Cult. coll. | Code | Species | Habitat -troph | Taxonomic classification | Raman identity | Cult. method |
| --- | --- | --- | --- | --- | --- | --- |
| priv |  | *Eimeria maxima* (A) (m) | E h p | SAR – Alveolata – **Apicomplexa** – Coccidia | Guanine | † |
| priv |  | *Psychodiella sergenti* | E h p | SAR – Alveolata – **Apicomplexa** – Gregarinasina | Guanine | – |
| NCMA | CCMP3155 | *Vitrella brassicaformis* | M a | SAR – Alveolata – **Chrompodellids** – Vitrellaceae | Guanine | – |
| ns |  | *Coleps* sp. | F h | SAR – Alveolata – **Ciliophora** – Prostomatea | Starch | – |
| ns |  | *Cyclidium* sp. | F h | SAR – Alveolata – **Ciliophora** – Oligohymenophorea | Guanine | – |
| ns |  | *Euplotes* sp. | F h | SAR – Alveolata – **Ciliophora** – Spirotrichea | Guanine | – |
| ns |  | *Euplotes* sp. | M h | SAR – Alveolata – **Ciliophora** – Spirotrichea | Guanine | – |
| ns |  | *Halteria* sp. | F h | SAR – Alveolata – **Ciliophora** – Spirotrichea | Guanine | – |
| ns |  | *Glaucoma* sp. | F h | SAR – Alveolata – **Ciliophora** – Oligohymenophorea | Guanine | – |
| ns |  | Hypotrichia gen. sp. | M h | SAR – Alveolata – **Ciliophora** – Spirotrichea | Guanine | – |
| ns |  | *Lembadion* sp. | M h | SAR – Alveolata – **Ciliophora** – Oligohymenophorea | Guanine | – |
| ns |  | *Oxytricha* gen. sp. | F h | SAR – Alveolata – **Ciliophora** – Spirotrichea | Guanine | – |
| ns |  | *Paramecium* sp. (C) (m) | F h | SAR – Alveolata – **Ciliophora** – Oligohymenophorea | Guanine | † |
| ns |  | Scuticulociliatia gen. sp. | M h | SAR – Alveolata – **Ciliophora** – Oligohymenophorea | Guanine | – |
| ns |  | *Vorticella* sp. | F h | SAR – Alveolata – **Ciliophora** – Oligohymenophorea | – | – |
| priv |  | *Borghiella* sp. | F a | SAR – Alveolata – **Dinoglagellata** – Suessiales | Guanine | – |
| NCMA | CCMP449 | *Heterocapsa triquetra* | M a | SAR – Alveolata – **Dinoglagellata** – Peridiniales | Guanine | f/2 |
| priv |  | *Glenodinium foliaceum* (B) | M a | SAR – Alveolata – **Dinoglagellata** – Peridiniales | Guanine | – |
| ns |  | *Gymnodinium* sp. | F a | SAR – Alveolata – **Dinoglagellata** – Gymnodiniales | Guanine | – |
| ns |  | *Peridinium* sp. | F a | SAR – Alveolata – **Dinoglagellata** – Peridiniales | Guanine | – |
| priv |  | *Sphaerodinium* sp. | F a | SAR – Alveolata – **Dinoglagellata** – Suessiales | Guanine | – |
| priv | 370 | *Symbiodinium microadriaticum* (m) | M a | SAR – Alveolata – **Dinoglagellata** – Suessiales | Guanine | f/2 |
| NCMA | CCMP829 | *Symbiodinium tridacnidorum* | M a | SAR – Alveolata – **Dinoglagellata** – Suessiales | Guanine | f/2 |
| ns |  | *Symbiodinium* sp.<br>isolated from *Capnella imbricata* | M a | SAR – Alveolata – **Dinoglagellata** – Suessiales | Guanine | f/2 |
| CAUP | J95 | *Achnanthidium* sp. | F a | SAR – Stramenopiles – **Ochrophyta** – **Bacillariophyceae** | Calcite | – |
| ns |  | *Asterionella formosa* | F a | SAR – Stramenopiles – **Ochrophyta** – **Bacillariophyceae** | – | – |
| ns |  | *Aulacoseira* sp. | F a | SAR – Stramenopiles – **Ochrophyta** – **Bacillariophyceae** | – | – |
| ns |  | *Encyonema* sp. | F a | SAR – Stramenopiles – **Ochrophyta** – **Bacillariophyceae** | Uric acid | – |
| ns |  | *Entomoneis ornata* | M a | SAR – Stramenopiles – **Ochrophyta** – **Bacillariophyceae** | – | – |
| ns |  | *Fragilaria* sp. | F a | SAR – Stramenopiles – **Ochrophyta** – **Bacillariophyceae** | Uric acid | – |
| ns |  | *Frustulia* sp. | F a | SAR – Stramenopiles – **Ochrophyta** – **Bacillariophyceae** | – | – |
| ns |  | *Gomphonema truncatum* | F a | SAR – Stramenopiles – **Ochrophyta** – **Bacillariophyceae** | – | – |

| Cult. coll. | Code | Species | Habitat -trophy | Taxonomic classification | Raman identity | Cult. method |
| --- | --- | --- | --- | --- | --- | --- |
| ns |  | *Licmophora* sp. | M a | SAR – Stramenopiles – **Ochrophyta** – **Bacillariophyceae** | Calcite | – |
| ns |  | *Melosira* sp. | F a | SAR – Stramenopiles – **Ochrophyta** – **Bacillariophyceae** | – | – |
| ns |  | *Navicula cryptocephala* | F a | SAR – Stramenopiles – **Ochrophyta** – **Bacillariophyceae** | Uric acid | – |
| ns |  | *Navicula* sp. | F a | SAR – Stramenopiles – **Ochrophyta** – **Bacillariophyceae** | Uric acid<br>Xanthine | – |
| ns |  | Naviculaceae gen. sp., *Seminavis*-like | M a | SAR – Stramenopiles – **Ochrophyta** – **Bacillariophyceae** | Guanine | – |
| ns |  | *Paralia sulcata* | M a | SAR – Stramenopiles – **Ochrophyta** – **Bacillariophyceae** | Calcite | – |
| CCAP | 1052/1B | *Phaeodactylum tricornutum* (m) | M a | SAR – Stramenopiles – **Ochrophyta** – **Bacillariophyceae** | Calcite | f/2 |
| ns |  | *Pinnularia viridis* | F a | SAR – Stramenopiles – **Ochrophyta** – **Bacillariophyceae** | – | – |
| ns |  | *Pleurosigma* sp. | M a | SAR – Stramenopiles – **Ochrophyta** – **Bacillariophyceae** | Uric acid | – |
| ns |  | *Stauroneis phoenicenteron* | F a | SAR – Stramenopiles – **Ochrophyta** – **Bacillariophyceae** | – | – |
| NCMA | CCMP1335 | *Thalassiosira pseudonana* (m) | M a | SAR – Stramenopiles – **Ochrophyta** – **Bacillariophyceae** | – | f/2 |
| ns |  | *Spumella* sp. | F a | SAR – Stramenopiles – **Ochrophyta** – **Synurophyceae** | Guanine | – |
| CAUP | B710 | *Synura hibernica* | F a | SAR – Stramenopiles – **Ochrophyta** – **Synurophyceae** | Guanine | – |
| CAUP | H4302 | *Eustigmatos polyphem* (b) | T a | SAR – Stramenopiles – **Ochrophyta** – **Eustigmatophyceae** | Guanine | BBM |
| NIES | 2146 | *Nannochloropsis oculata* (K) (b) (m) | M a | SAR – Stramenopiles – **Ochrophyta** – **Eustigmatophyceae** | Guanine | f/2 |
| NIES | 2860 | *Vacuoliviride crystalliferum*<sup>#</sup> | Un a | SAR – Stramenopiles – **Ochrophyta** – **Eustigmatophyceae** | Guanine<br>Chrysolaminarin | BBM |
| ns |  | *Cystoseira* sp. | M a | SAR – Stramenopiles – **Ochrophyta** – **Phaeophyceae** | Calcite | – |
| ns |  | *Dictyota dichotoma* | M a | SAR – Stramenopiles – **Ochrophyta** – **Phaeophyceae** | Calcite | – |
| ns |  | *Padina pavonica* | M a | SAR – Stramenopiles – **Ochrophyta** – **Phaeophyceae** | Calcite | – |
| ns |  | *Actinophrys* sp. | F a | SAR – Stramenopiles – **Ochrophyta** – **Actinophryidae** | Guanine | – |
| ns |  | *Gonyostomum* sp. | F a | SAR – Stramenopiles – **Ochrophyta** – **Raphidophyceae** | Guanine | – |
| CAUP | D301 | *Botrydiopsis intercedens* | F a | SAR – Stramenopiles – **Ochrophyta** – **Xanthophyceae** | Guanine | BBM |
| UTEX | B 2999 | *Heterococcus* sp. | F a | SAR – Stramenopiles – **Ochrophyta** – **Xanthophyceae** | – | BBM |
| ns |  | *Ophiocytium* sp. | F a | SAR – Stramenopiles – **Ochrophyta** – **Xanthophyceae** | Guanine | – |
| CCALA | 517 | *Tribonema aequale* (F) (m) | F a | SAR – Stramenopiles – **Ochrophyta** – **Xanthophyceae** | Guanine | BBM |
| ns |  | *Xanthonema* sp. | F a | SAR – Stramenopiles – **Ochrophyta** – **Xanthophyceae** | Guanine | – |
| DBM | CO3I | *Schizochytrium* sp. (b) (m) | M h | SAR – Stramenopiles – **Labyrinthulomycota** | Guanine | ASW-L |
| priv |  | *Cafeteria roenbergensis* | M h | SAR – Stramenopiles – **Bicosoecida** | – | f/2* |
| priv |  | *Cafileria marina* | M h | SAR – Stramenopiles – **Bicosoecida** | Guanine | f/2* |
| CCF | 3762 | *Phytophthora cactorum* | E h | SAR – Stramenopiles – **Oomycota** | – | PDA |
| CCF | 4738 | *Phytophthora rosacearum*-like | E h | SAR – Stramenopiles – **Oomycota** | – | PDA |
| priv | Ther | *Blastocystis* sp. isolated from *Testudo hermanni* | E h | SAR – Stramenopiles – **Opalinata** – Blastocystae | – | TYM |
| NCMA | CCMP2755 | *Bigelowiella natans* (L) (m) | M a | SAR – Rhizaria – **Cercozoa** – **Chlorarachnea** | Guanine | f/2 |
| priv | LITO | *Cercomonadida* gen. sp. | F h | SAR – Rhizaria – **Cercozoa** – **Paracercomonadida** | Guanine | Fw |
| ns |  | *Viridiraptor invadens* | F h | SAR – Rhizaria – **Cercozoa** – **Viridiraptoridae** | Guanine | – |
| ns |  | *Euglypha* sp. | F h | SAR – Rhizaria – **Cercozoa** – **Euglyphida** | Guanine | – |
| ns |  | *Trinema* sp. | F h | SAR – Rhizaria – **Cercozoa** – **Euglyphida** | Guanine | – |
| ns |  | *Marinomyxa marina* | M h | SAR – Rhizaria – Endomyxa – **Phytomyxea** | – | – |
| ns |  | *Acantharea* gen. sp. | M h | SAR – Rhizaria – **Retaria** – Acantharea | Celestite | † |
| ns |  | *Globigerinidae* gen. sp. | M h | SAR – Rhizaria – **Retaria** – Foramanifera | – | † |
| ns |  | *Globorotaliidae* gen. sp. | M h | SAR – Rhizaria – **Retaria** – Foramanifera | Calcite | † |
| priv | LIS-Tel | *Telonema* sp. | M h | **Telonemida** | Guanine | f/2B* |
| NCMA | CCMP371 | *Emiliana huxleyi* (m) | M a | Haptista – **Haptophyta** | Guanine | f/2* |
| RCC | RCC1350 | *Isochrysis* sp. (M) (b) | M a | Haptista – **Haptophyta** | Xanthine<br>Uric acid | f/2* |
| ns |  | *Raphidiophrys* sp. | F h | Haptista – **Centroplasthelida** | Guanine | – |
| CAUP | F105 | *Cryptomonas* sp. (O) | F a | Cryptista – **Cryptophyta** | Uric acid | BBM |
| ns |  | *Chroomonas* sp. | F a | Cryptista – **Cryptophyta** | Uric acid<br>Guanine<br>monohydrate | – |
| CAUP | O101 | *Glaucocystis nostochinearum* | F a | Archaeplastida – **Glaucophyta** | Xanthine | BBM |
| ns |  | *Asparagopsis taxiformis* | M a | Archaeplastida – **Rhodophyta** – **Florideophyceae** | Calcite | – |
| CAUP | L201 | *Asterocytis ramosa* | M a | Archaeplastida – **Rhodophyta** – **Bangiophyceae** | Carotenoids,<br>lipids | f/2 |
| CCALA | 971 | *Audouinella* sp. | M a | Archaeplastida – **Rhodophyta** – **Florideophyceae** | – | f/2 |
| ns |  | *Ceramium* sp. | M a | Archaeplastida – **Rhodophyta** – **Florideophyceae** | Calcite | – |
| ns |  | *Hildebrandia rivularis* | M a | Archaeplastida – **Rhodophyta** – **Florideophyceae** | – | – |
| CCALA | 416 | *Porphyridium purpureum* (b) | M a | Archaeplastida – **Rhodophyta** – **Porphyridiophyceae** | – | f/2 |
| CCALA | 925 | *Rhodella violacea* | M a | Archaeplastida – **Rhodophyta** – **Rhodellophyceae** | Lipids | f/2 |
| ns |  | *Asterococcus* sp. | F a | Archaeplastida – Viridiplantae – Chlorophyta – **Chlorophyceae** | Guanine | – |
| ns |  | *Bulbochaete* sp. | F a | Archaeplastida – Viridiplantae – Chlorophyta – **Chlorophyceae** | Guanine | – |
| CRC | CC-1690 | *Chlamydomonas reinhardtii*<sup>#</sup> (m) | F a | Archaeplastida – Viridiplantae – Chlorophyta – **Chlorophyceae** | Guanine | BBM |
| CAUP | G224 | *Chlamydomonas geitleri* | F a | Archaeplastida – Viridiplantae – Chlorophyta – **Chlorophyceae** | Guanine | BBM |
| ns |  | *Chlamydomonas* sp. | F a | Archaeplastida – Viridiplantae – Chlorophyta – **Chlorophyceae** | Guanine | – |
| priv | AMAZONIE | *Chlamydomonadales* gen. sp. | F h | Archaeplastida – Viridiplantae – Chlorophyta – **Chlorophyceae** | Guanine | Fw |
| priv | KBEL1C | *Polytoma* sp. | F h | Archaeplastida – Viridiplantae – Chlorophyta – **Chlorophyceae** | Guanine | Fw |
| CAUP | H 6902 | *Chlorochytrium lemnae* | E – F a | Archaeplastida – Viridiplantae – Chlorophyta – **Chlorophyceae** | Guanine | BBM |
| priv |  | *Chloromonas arctica* | S a | Archaeplastida – Viridiplantae – Chlorophyta – **Chlorophyceae** | Guanine | BBM |
| ns |  | *Desmodesmus quadricauda* (m) | F a | Archaeplastida – Viridiplantae – Chlorophyta – **Chlorophyceae** | Guanine | – |
| ns |  | *Desmodesmus* sp. | F a | Archaeplastida – Viridiplantae – Chlorophyta – **Chlorophyceae** | Guanine | – |
| ns |  | *Eudorina* sp. | F a | Archaeplastida – Viridiplantae – Chlorophyta – **Chlorophyceae** | Guanine | – |
| ns |  | *Gleocystis* sp. | F a | Archaeplastida – Viridiplantae – Chlorophyta – **Chlorophyceae** | Guanine | – |
| ns |  | *Microspora* sp. | F a | Archaeplastida – Viridiplantae – Chlorophyta – **Chlorophyceae** | Guanine | – |
| ns |  | *Monactinus simplex* | F a | Archaeplastida – Viridiplantae – Chlorophyta – **Chlorophyceae** | Guanine | – |
| CAUP | H2908 | *Monoraphidium contortum* (b) | F a | Archaeplastida – Viridiplantae – Chlorophyta – **Chlorophyceae** | Guanine | BBM |
| ns |  | *Oedogonium* sp. | F a | Archaeplastida – Viridiplantae – Chlorophyta – **Chlorophyceae** | Guanine | – |
| ns |  | *Pediastrum boryanum* | F a | Archaeplastida – Viridiplantae – Chlorophyta – **Chlorophyceae** | Guanine | – |
| CAUP | H2308 | *Pediastrum duplex* (I) | F a | Archaeplastida – Viridiplantae – Chlorophyta – **Chlorophyceae** | Guanine | BBM |
| ns |  | *Tetraedron* sp. | F a | Archaeplastida – Viridiplantae – Chlorophyta – **Chlorophyceae** | Guanine | – |
| ns |  | *Tetraspora* sp. | F a | Archaeplastida – Viridiplantae – Chlorophyta – **Chlorophyceae** | Guanine | – |
| CAUP | H 1917 | *Chlorella vulgaris* (N) (b) (m) | F a | Archaeplastida – Viridiplantae – Chlorophyta – **Trebouxiophyceae** | Xanthine | BBM* |
| CAUP | H 5107 | *Coccomyxa elongata* | T a | Archaeplastida – Viridiplantae – Chlorophyta – **Trebouxiophyceae** | Guanine | BBM |
| ns |  | *Dictyosphaerium* sp. (b) | F a | Archaeplastida – Viridiplantae – Chlorophyta – **Trebouxiophyceae** | Uric acid | – |
| ns |  | *Keratococcus* sp. | F a | Archaeplastida – Viridiplantae – Chlorophyta – **Trebouxiophyceae** | Guanine | – |
| ns |  | *Micractinium* sp. | F a | Archaeplastida – Viridiplantae – Chlorophyta – **Trebouxiophyceae** | Guanine | – |
| CAUP | H8801 | *Phyllosiphon arisari* | F a p | Archaeplastida – Viridiplantae – Chlorophyta – **Trebouxiophyceae** | Lipids | BBM |
| CAUP | H8605 | *Symbiochloris tschermakiae* | E/T a | Archaeplastida – Viridiplantae – Chlorophyta – **Trebouxiophyceae** | Lipids | BBM |
| ns |  | *Anadyomene* sp. | M a | Archaeplastida – Viridiplantae – Chlorophyta – **Ulvophyceae** | Calcite | – |
| ns |  | *Cladophora* sp. | F a | Archaeplastida – Viridiplantae – Chlorophyta – **Ulvophyceae** | Guanine | – |
| ns |  | *Cladophora* sp. | M a | Archaeplastida – Viridiplantae – Chlorophyta – **Ulvophyceae** | Calcite | – |
| ns |  | *Enteromorpha* sp. | M a | Archaeplastida – Viridiplantae – Chlorophyta – **Ulvophyceae** | Calcite | – |
| CAUP | H5301 | *Scotinosphaera gibberosa* | F/T a | Archaeplastida – Viridiplantae – Chlorophyta – **Ulvophyceae** | Carotenoids, lipids | BBM |
| CAUP | J1601 | *Trentepohlia* sp. | T a | Archaeplastida – Viridiplantae – Chlorophyta – **Ulvophyceae** | Carotenoids, lipids | BBM |
| CAUP | M101 | *Prasinocladus ascus* | M a | Archaeplastida – Viridiplantae – Chlorophyta – **Chlorodendrophyceae** | Guanine monohydrate<br>Uric acid | – |
| CAUP | M201 | *Tetraselmis subcordiformis* (H) (b) | M a | Archaeplastida – Viridiplantae – Chlorophyta – **Chlorodendrophyceae** | Guanine monohydrate<br>Uric acid | f/2 |
| RCC | RCC745 | *Ostreococcus tauri* (m) | M a | Archaeplastida – Viridiplantae – Chlorophyta – **Mamiellophyceae** | – | f/2 |
| ns |  | *Nephroselmis* sp. | M a | Archaeplastida – Viridiplantae – Chlorophyta – **Nephrophyceae** | Guanine | – |
| ns |  | *Pyramimonas* sp. | M a | Archaeplastida – Viridiplantae – Chlorophyta – **Pyramimonadophyceae** | Guanine | – |
| NIES | 995 | *Mesostigma viride* | F a | Archaeplastida – Viridiplantae – Streptophyta – **Mesostigmatophyta** | – | – |
| CAUP | H7601 | *Chlorokybus atmophyticus* | T a | Archaeplastida – Viridiplantae – Streptophyta – **Chlorokybophyceae** | Guanine | BBM |
| NIES | 2285 | *Klebsormidium flaccidum*<sup>#</sup> (P) | T a | Archaeplastida – Viridiplantae – Streptophyta – **Klebsormidiophyceae** | Uric acid | BBM |
| ns |  | *Coleochaete* sp. | F a | Archaeplastida – Viridiplantae – Streptophyta – **Coleochaetophyceae** | Calcium oxalate | – |
| NIES | 1604 | *Chara braunii* | F a | Archaeplastida – Viridiplantae – Streptophyta – **Charophyceae** | – | – |
| CAUP | K 801 | *Actinotaenium* sp. | F a | Archaeplastida – Viridiplantae – Streptophyta – **Zygnematophyceae** | Uric acid | BBM |
| NIES | 67 | *Closterium peracerosum-strigosum-littorale* complex (m) | F a | Archaeplastida – Viridiplantae – Streptophyta – **Zygnematophyceae** | Baryte<br>Starch | BBM |
| ns |  | *Cosmarium* sp. | F a | Archaeplastida – Viridiplantae – Streptophyta – **Zygnematophyceae** | Uric acid<br>Baryte | – |
| ns |  | *Cylindrocystis* sp. | T a | Archaeplastida – Viridiplantae – Streptophyta – **Zygnematophyceae** | Calcium oxalate | – |
| ns |  | *Euastrum humerosum* | F a | Archaeplastida – Viridiplantae – Streptophyta – **Zygnematophyceae** | Uric acid | – |
| CAUP | K101 | *Mesotaenium caldariorum* | F a | Archaeplastida – Viridiplantae – Streptophyta – **Zygnematophyceae** | Uric acid<br>Guanine monohydrate | BBM* |
| ns |  | *Mougeotia* sp. | F a | Archaeplastida – Viridiplantae – Streptophyta – **Zygnematophyceae** | Uric acid | – |
| ns |  | *Netrium* sp. | F a | Archaeplastida – Viridiplantae – Streptophyta – **Zygnematophyceae** | Uric acid | – |
| NIES | 303 | *Penium margaritaceum* (Q) | F a | Archaeplastida – Viridiplantae – Streptophyta – **Zygnematophyceae** | Uric acid | BBM |
| ns |  | *Spirogyra* sp. | F a | Archaeplastida – Viridiplantae – Streptophyta – **Zygnematophyceae** | Baryte | – |
| ns |  | *Staurostrum* sp. | F a | Archaeplastida – Viridiplantae – Streptophyta – **Zygnematophyceae** | Uric acid | – |
| ns |  | *Tetmemorus* sp. | F a | Archaeplastida – Viridiplantae – Streptophyta – **Zygnematophyceae** | Uric acid | – |
| ns |  | *Xanthidium* sp. | F a | Archaeplastida – Viridiplantae – Streptophyta – **Zygnematophyceae** | Uric acid | – |
| ns |  | *Halophila stipulacea* | M a | Archaeplastida – Viridiplantae – Streptophyta – **Embryophyta** | Calcium oxalate monohydrate,<br>dihydrate | – |
| priv | BY-2 | *Nicotiana tabacum* | T a | Archaeplastida – Viridiplantae – Streptophyta – **Embryophyta** | Calcium oxalate monohydrate | MS-1 |
| priv |  | *Physcomitrela patens* | T a | Archaeplastida – Viridiplantae – Streptophyta – **Embryophyta** | Calcium oxalate monohydrate | MS-2 |
| priv | BW2 | *Hemimastix kukwesjijk* | F h | **Hemimastigophora** | Guanine | FwB2* |
| ns |  | *Entosiphon* sp. | F a | Excavata – Discoba – **Euglenozoa – Euglenida** | Guanine | – |
| ns |  | *Euglena* sp. (m) | F a | Excavata – Discoba – **Euglenozoa – Euglenida** | Guanine | – |
| NCMA | CCMP1594 | *Eutreptiella gymnastica* (D) (m) | M a | Excavata – Discoba – **Euglenozoa – Euglenida** | Guanine | f/2 |
| ns |  | *Lepocynclis oxyuris* | F a | Excavata – Discoba – **Euglenozoa – Euglenida** | Guanine | – |
| ns |  | *Monomorphina pyrum* | F a | Excavata – Discoba – **Euglenozoa – Euglenida** | Guanine | – |
| ns |  | *Phacus* sp. | F a | Excavata – Discoba – **Euglenozoa – Euglenida** | Guanine | – |
| priv | SVATBA | *Rhabdomonas* sp. | F h | Excavata – Discoba – **Euglenozoa – Euglenida** | Guanine | Fw |
| ns |  | *Trachelomonas* sp. | F a | Excavata – Discoba – **Euglenozoa – Euglenida** | Guanine | – |
| priv | RIOZ | Free-living Kinetoplastea gen. sp. | F h | Excavata – Discoba – **Euglenozoa – Kinetoplastea** | Guanine | Fw |
| ns |  | *Bodo* sp. | F h | Excavata – Discoba – **Euglenozoa – Kinetoplastea** | Guanine | – |
| priv | M09 | '*jaculum*' gen. sp. | E h p | Excavata – Discoba – **Euglenozoa – Kinetoplastea** | – | BHI |
| priv |  | *Trypanosoma mega* | E h p | Excavata – Discoba – **Euglenozoa – Kinetoplastea** | – | BHI |
| priv | YPF 1621 | *Namystynia karyoxenos* | M h | Excavata – Discoba – **Euglenozoa – Diplonemea** | Guanine | ASW-D |
| priv | 1.7 clone | *Rhynchopus* sp. | M h | Excavata – Discoba – **Euglenozoa – Diplonemea** | Guanine | ASW-D |
| priv | DT1610 | *Flectonema* sp. | M h | Excavata – Discoba – **Euglenozoa – Diplonemea** | Guanine monohydrate | ASW-D |
| priv | RUM4AN | *Psalteriomonas lanterna* | F h | Excavata – Discoba – **Heterolobosea** | Guanine | Fw |
| priv | NEG-M | *Naegleria gruberi* (E) (m) | F h | Excavata – Discoba – **Heterolobosea** | Guanine | M7* |
| priv | BUSSPRAND | *Velundella trypanoides* | M h | Excavata – Discoba – **Jakobida** | – | JM |
| priv | AND | *Stygiella agilis* | M h | Excavata – Discoba – **Jakobida** | – | JM |
| priv | SPINDL2 | *Gyromonas ambulans* | F h | Excavata – **Metamonada** – Fornicata | Guanine | Fw* |
| priv | TUN2 | *Hexamita* sp. | F h | Excavata – **Metamonada** – Fornicata | Guanine | Fw* |
| priv | LITO | *Trepomonas rotans* | F h | Excavata – **Metamonada** – Fornicata | Guanine | Fw* |
| priv | BREZ2C | *Trepomonas* sp. | F h | Excavata – **Metamonada** – Fornicata | Guanine | Fw* |
| priv | CONGO | *Trepomonas steinii* | F h | Excavata – **Metamonada** – Fornicata | Guanine | LB |
| priv | Ncub | *Macrotrichomonoides* sp. isolated from *Neotermes cubanus* | E h | Excavata – **Metamonada** – Parabasalia | Guanine | – |
| priv | 249 | *Gefionella okellyi* (J) | F h | **Malawimonadida** | Guanine | f/2B* |
| ATCC | HM-1:IMSS | *Entamoeba histolytica* | E h | Amoebozoa – Evosea – **Archamoebae** | Lipids | TYI* |
| ATCC | 30984 | *Mastigamoeba balamuthi* | F h | Amoebozoa – Evosea – **Archamoebae** | Xanthine<br>Lipids | PYGC* |
| priv | GO7 | *Mastigella eilhardi* | F h | Amoebozoa – Evosea – **Archamoebae** | Guanine | Fw* |
| ns |  | *Fuligo septica* | T h | Amoebozoa – Evosea – **Eumycetozoa** | Guanine | – |
| priv |  | *Physarum polycephalum* | T h | Amoebozoa – Evosea – **Eumycetozoa** | Xanthine | OG * |
| priv | Neff | *Acanthamoeba castellanii* | F h | Amoebozoa – **Discosea** – Centramoebia | Xanthine<br>Lipids | Neff* |
| ns |  | *Mayorella* sp. | F h | Amoebozoa – **Discosea** – Flabellinia | Xanthine | – |
| ns |  | *Mayorella* sp. | M h | Amoebozoa – **Discosea** – Flabellinia | Xanthine | – |
| priv | WFP252 | *Neoparamoeba* sp. | M h | Amoebozoa – **Discosea** – Flabellinia | – | MAM |
| ns |  | *Thecamoeba* sp. | F h | Amoebozoa – **Discosea** – Flabellinia | Guanine | – |
| ns |  | *Vannella* sp. | F h | Amoebozoa – **Discosea** – Flabellinia | Guanine | – |
| ns |  | *Diffugia* sp. | F h | Amoebozoa – **Tubulinea** – Arcellinida | Guanine | – |
| ns |  | *Rhizamoeba* sp. | F h | Amoebozoa – **Tubulinea** – Leptomyxida | – | – |
| ns |  | *Saccamoeba* sp. | F h | Amoebozoa – **Tubulinea** – Hartmannellidae | Unidentified | – |
| priv | MSEDG | *Saccamoeba* sp. | F h | Amoebozoa – **Tubulinea** – Hartmannellidae | Unidentified | FwAM* |
| priv | 4391/l | *Vermamoeba vermiformis* | F h | Amoebozoa – **Tubulinea** – Echinamoebida | – | FwAM* |
| priv | LIS-Man | *Mantamonas* sp. | M h | CRuMs – **Mantamonadida** | – | f/2B* |
| priv | FB10 | *Subulatomonas* sp. | M h | Obazoa – **Breviatea** | – | LBM |
| priv | C3C | Choanoflagellata gen. sp. | Hp h | Obazoa – Opisthokonta – **Choanoflagellata** | – | HM-V* |
| ns |  | Chaetonotidae gen. sp. | F h | Obazoa – Opisthokonta – **Metazoa** – Gastrotricha | Guanine | – |
| ns |  | Nematoda gen. sp. | F/T h | Obazoa – Opisthokonta – **Metazoa** – Nematoda | Uric acid | – |
| ns |  | *Oncorhynchus mykiss* (scale) | F h | Obazoa – Opisthokonta – **Metazoa** – Chordata | Guanine | – |
| CCF | 2912 | *Aspergillus nidulans* | F/T h | Obazoa – Opisthokonta – **Fungi** – Ascomycota – Eurotiales | – | MEA |
| ATCC | SC5314 | *Candida albicans* | F/T h | Obazoa – Opisthokonta – **Fungi** – Ascomycota – Saccharomycetales | Guanine | MEA |
| EUROS<br>CARF | BY4741 | *Saccharomyces cerevisiae* | T h | Obazoa – Opisthokonta – **Fungi** – Ascomycota – Saccharomycetales | – | YPAD |
| CCF | 3485 | *Neurospora sitophila* | F/T h | Obazoa – Opisthokonta – **Fungi** – Ascomycota – Sordariales | – | MEA |
| CCVC | CQ1 | *Nosema bombycis* | E h p | Obazoa – Opisthokonta – **Opisthosporidia** – **Microsporidia** | – | – |
| ns |  | *Woronichinia naegeliana* | F a | Eubacteria – Cyanobacteria | Aerotopes | – |
| ns |  | *Oscillatoria* sp. | F a | Eubacteria – Cyanobacteria | Carotenoids,<br>lipids | – |
| priv |  | *Bacillus subtilis* | T h | Eubacteria – Firmicutes | – | – |

**Table S2.** Overview of the Concentrative nucleoside transporter (CNT) distribution among eukaryotes based on HMM search in the database of 57 eukaryotic genomes.

| **Taxonomic rank** | **Presence of CNT** |
| --- | --- |
| Amoebozoa | yes |
| Holomycota | yes |
| Chrompodellids | yes |
| Streptophyta | no |
| Rhizaria | yes |
| Stramenopiles | no |
| Holozoa | yes |
| Chlorophyta | no |
| Haptophyta | yes |
| Cryptophyceae | yes |
| Apicomplexa | no |
| Rhodophyta | no |
| Euglenozoa | no |
| Metamonada | no |
| Ciliophora | no |
| Heterolobosea | no |
| Perkinsidae | no |
| Apusomonadida | no |

**Table S3.** Overview of the Equilibrative nucleoside transporter (ENT) distribution among eukaryotes based on HMM search in the database of 57 eukaryotic genomes.

| **Taxonomic rank** | **Presence of ENT** |
| --- | --- |
| Amoebozoa: | yes |
| Holomycota: | yes |
| Chrompodellids | yes |
| Streptophyta | yes |
| Rhizaria | yes |
| Stramenopiles | yes |
| Holozoa | yes |
| Chlorophyta | yes |
| Haptophyta | yes |
| Cryptophyceae | yes |
| Apicomplexa | yes |
| Rhodophyta | yes |
| Euglenozoa | yes |
| Metamonada | yes |
| Ciliophora | yes |
| Heterolobosea | yes |
| Perkinsidae | yes |
| Apusomonadida | yes |

**Movie S1.**

Polarized light microscopy of crystalline inclusions in SAR. https://youtu.be/cMkMJthq5KQ

**Movie S2.**

Polarized light microscopy of crystalline inclusions in Haptista, Cryptista and Telonemia. https://youtu.be/Z30CDbWqOhc

**Movie S3.**

Polarized light microscopy of crystalline inclusions in Archaeplastida. https://youtu.be/2ZXfOdpsJcU

**Movie S4.**

Polarized light microscopy of crystalline inclusions in Amoebozoa. https://youtu.be/DUbA5e_1_BE

**Movie S5.**

Polarized light microscopy of crystalline inclusions in Opisthokonta. https://youtu.be/kyEzbo-IbiM

**Movie S6.**

Polarized light microscopy of crystalline inclusions in Excavata. https://youtu.be/XWzNBLmE01A

**Movie S7.**

Polarized light microscopy of crystalline inclusions in Hemimastigophora. https://youtu.be/gnUZhZRRfcw

**Movie S8.**

Polarized light microscopy of crystalline inclusions in Prokaryota – Cyanobacteria. https://youtu.be/8yGo161rdJU

**Correspondence and requests for materials should be addressed to:**

